# POGZ modulates the DNA damage response in a HP1-dependent manner

**DOI:** 10.1101/2021.06.28.447216

**Authors:** John Heath, Estelle Simo Cheyou, Steven Findlay, Vincent M Luo, Edgar Pinedo Carpio, Jeesan Lee, Billel Djerir, Xiaoru Chen, Théo Morin, Benjamin Lebeau, Martin Karam, Halil Bagci, Damien Grapton, Josie Ursini-Siegel, Jean-Francois Côté, Michael Witcher, Stéphane Richard, Alexandre Maréchal, Alexandre Orthwein

**Author notes:** Institute of Biochemistry, ETH Zürich, Zürich, Switzerland. Address correspondence to: Alexandre Orthwein M.Sc., Ph.D. Lady Davis Institute, Segal Cancer Centre Jewish General Hospital, Room E-447 3755 Chemin de la Côte-Sainte-Catherine Montreal, Quebec, H3T 1E2 Canada Tel: ext. 24252.

## Abstract

The heterochromatin protein HP1 plays a central role in the maintenance of genome stability, in particular by promoting homologous recombination (HR)-mediated DNA repair. However, little is still known about how HP1 is controlled during this process. Here, we describe a novel function of the POGO transposable element derived with ZNF domain protein (POGZ) in the regulation of HP1 during the DNA damage response *in vitro*. POGZ depletion delays the resolution of DNA double-strand breaks (DSBs) and correlates with an increased sensitivity to different DNA damaging agents, including the clinically-relevant Cisplatin and Talazoparib. Mechanistically, POGZ promotes homology-directed DNA repair pathways by retaining the BRCA1/BARD1 complex at DSBs, in a HP1-dependent manner. *In vivo* CRISPR inactivation of *Pogz* is embryonically lethal and *Pogz* haplo-insufficiency (*Pogz*^+/Δ^) results in a developmental delay, impaired intellectual abilities, a hyperactive behaviour as well as a compromised humoral immune response in mice, recapitulating the main clinical features of the White Sutton syndrome (WHSUS). Importantly, *Pogz*^+/Δ^ mice are radiosensitive and accumulate DSBs in diverse tissues, including the spleen and the brain. Altogether, our findings identify POGZ as an important player in homology-directed DNA repair both *in vitro* and *in vivo,* with clinical implications for the WHSUS.

## INTRODUCTION

DNA double-strand breaks (DSBs) are among the most cytotoxic DNA lesions, in part due to their highly recombinogenic and pro-apoptotic potential (Iyama & Wilson, 2013). Inaccurate resolution of DSBs can result in gross genomic rearrangements that drive genomic instability, a characteristic feature of several human genetic disorders and cancer subtypes (Bunting & Nussenzweig, 2013). To avoid this deleterious outcome, cells have deployed a complex network of proteins that detect and signal these DNA lesions for subsequent repair (Polo & Jackson, 2011). Ultimately, two major pathways can be mobilized to repair these DNA lesions: non-homologous end-joining (NHEJ) and homologous recombination (HR) (Hustedt & Durocher, 2016). While NHEJ is active throughout the cell cycle, HR is only active in the S/G2 phases, as it requires a sister chromatid as a template for the faithful repair of these DNA lesions. Importantly, several additional elements influence DNA repair pathway choice, including the complexity of the DNA ends and the epigenetic context (Scully, Panday et al., 2019).

One key step in the “decision-making” process underpinning DNA repair pathway choice relies on the antagonism between 53BP1 and BRCA1 (Densham & Morris, 2019). On one hand, 53BP1 relies on both H4K20me^2^/me^3^ and RNF168-mediated ubiquitination of H2A variants for its recruitment to DSBs (Botuyan, Lee et al., 2006, Fradet-Turcotte, Canny et al., 2013, Huyen, Zgheib et al., 2004). On the other hand, BRCA1, which exists as an obligate heterodimer with BARD1, is known to be rapidly recruited to DNA damage sites in a poly-ADP-ribose (PAR)-and ATM-dependent manner (Li & Yu, 2013, Manke, Lowery et al., 2003, Yu, Chini et al., 2003). Importantly, the identification of BARD1 as a reader of unmethylated histone H4 at lysine 20 (H4K20me^0^) (Nakamura, Saredi et al., 2019), a post-replicative chromatin mark, shed new light on how the BRCA1/BARD1 complex counteracts 53BP1 at sites of DNA damage in the S/G2 phases of the cell cycle (Chapman, Barral et al., 2013, Chapman, Sossick et al., 2012, Escribano-Diaz & Durocher, 2013, Feng, Fong et al., 2013, Zimmermann, Lottersberger et al., 2013), thereby initiating DNA end resection and favoring homology-directed DNA repair.

Several additional factors can influence the localization of the BRCA1/BARD1 complex (Greenberg, Sobhian et al., 2006). For instance, the rapid accumulation of the heterochromatin protein HP1 at DNA damage sites (Ayoub, Jeyasekharan et al., 2008, Ayoub, Jeyasekharan et al., 2009, Baldeyron, Soria et al., 2011, Luijsterburg, Dinant et al., 2009, Zarebski, Wiernasz et al., 2009), and the specific recognition of its γ isoform (HP1-γ) by BARD1 through a consensus PxVxL motif (Wu et al., 2015), have emerged as key pre-requisites for the presence of BRCA1 at DSBs (Lee, Kuo et al., 2013), and the commitment towards homology-directed DNA repair (Wu, Nishikawa et al., 2015, Wu, Togashi et al., 2016). Interestingly, the mobilization of HP1 correlates with dynamic waves of H3K9 methylation and demethylation around these DNA lesions (Ayrapetov, Gursoy-Yuzugullu et al., 2014, Falk, Lukasova et al., 2007, Fnu, Williamson et al., 2011, Young, McDonald et al., 2013), pointing towards a methylation-dependent mobilization of the HP1-BARD1-BRCA1 axis (Wu et al., 2015). However, conflicting reports have suggested that HP1 could accumulate at DSBs by interacting with local DNA repair factors through its chromoshadow (CSD) domain (Luijsterburg et al., 2009, Zarebski et al., 2009), thereby eliminating H3K9 methylation as a pre-requisite for the mobilization of the HP1-BARD1-BRCA1 axis. In both models, little is known about how this multiprotein complex is regulated during the DNA damage response and whether any additional factor may control the mobilization of the BRCA1/BARD1 complex to DSBs.

We therefore sought to gain further insight into the contribution of HP1 during genotoxic stress by using a biotin-based labelling approach, called BioID (Roux, Kim et al., 2012). As mammals encode three distinct isoforms (-α, -β, and -γ), we mapped their respective proximal interactomes under normal and genotoxic stress conditions and identified the POGO transposable element derived with ZNF domain protein (POGZ) as one of the most abundant interactors of the different HP1 isoforms in both conditions, as previously described (Vermeulen, Eberl et al., 2010). POGZ has been implicated in the dissociation of HP1-α from mitotic chromosomes, thereby influencing their proper segregation (Nozawa, Nagao et al., 2010). Still, its role in the maintenance of genome stability during interphase remains elusive (Baude, Aaes et al., 2016), prompting us to investigate its contribution to the DNA damage response. Interestingly, POGZ depletion delays the resolution of DSBs in S/G2 phases of the cell cycle, which correlates with a prolonged G2 DNA damage checkpoint. Using well-established GFP-based DNA repair assays, we established that POGZ regulates both HR and single-strand annealing (SSA) pathways. Importantly, we noted that POGZ is rapidly recruited to DNA damage sites where it allows the presence of both BRCA1 and BARD1 at DSBs in a HP1-dependent manner. Subsequent *in vivo* analysis demonstrated the critical role of Pogz in murine embryogenesis and *Pogz* haplo-insufficient (*Pogz*^+/Δ^) mice display a significant growth defect, a deficit in intellectual abilities, a hyperactive behaviour as well as a compromised humoral immune response, recapitulating the main clinical features of the White Sutton syndrome (WHSUS) (Dentici, Niceta et al., 2017, Ferretti, Barresi et al., 2019, Fukai, Hiraki et al., 2015, Matsumura, Nakazawa et al., 2016, Samanta, Ramakrishnaiah et al., 2020, Tan, Zou et al., 2016, Ye, Cho et al., 2015, Zhao, Quan et al., 2019a, Zhao, Tan et al., 2019b). Strikingly, *Pogz*^+/Δ^ mice are hypersensitive to ionizing radiation (IR) and present constitutive DSBs in several tissues. Furthermore, *Pogz*^+/Δ^ B-cells have impaired survival capacities following the induction of programmed DSBs *ex vivo*. Altogether, our data describe a novel role of POGZ in the regulation of homology-directed DNA repair pathways through the HP1-BRCA1-BARD1 axis, with new perspectives for the aetiology of the WHSUS.

## RESULTS

### Identification of POGZ as a high confidence interactor of the HP1 isoforms

To gain better insight into the contribution of the different HP1 isoforms during the DNA damage response (Fig.1A), we used the BioID labelling technique, which allows the monitoring of proximal/transient interactions (Expanded View Fig.1A) (Varnaite & MacNeill, 2016). Briefly, HP1 proteins were fused to a mutant of an *E. coli* biotin-conjugating enzyme (BirA*) and stably expressed in the human embryonic kidney 293 (HEK293) cell line using the Flp-In/T-REX system and validated for their ability to biotinylate proteins that come in close proximity or directly interact with them (Expanded View Fig.1B). Labelled proteins were subsequently purified by streptavidin affinity and identified by mass spectrometry. This approach was carried out in cells exposed to the radiomimetic DNA damaging drug neocarzinostatin (NCS) or the solvent control (Ctrl). For each HP1 isoform, we identified ∼500 different high-confidence interactors that were common to both experimental conditions (Fig.1B and Appendix Table S1). Gene set enrichment analysis (GSEA) demonstrated that proteins involved in cell cycle progression, in particular during M phase, epigenetic regulation of gene expression and DNA repair were commonly found in the interactome of HP1-α, -β and –γ in both Ctrl and NCS-treated conditions (Fig.1C, Expanded View Fig.1D and Appendix Table S2). As expected, histone variants (HIST1H2BL, HIST3H2BB, H2AFY, H2AFZ), chromatin-associated proteins (CHAF1A, HMGN2), and histone methyltransferases (EHMT1, WIZ) emerged as the most abundant interactors of the different HP1 isoforms in both Ctrl and NCS-treated conditions (Fig.1D). Additionally, we identified POGZ as a high-confidence interactor of the three HP1 isoforms (Fig.1D), which tends to be more biotinylated upon NCS treatment compared to Ctrl conditions (Appendix Table S1), particularly with the BirA-HP1-γ construct. This prompted us to investigate whether POGZ functions during the DNA damage response.

**Figure 1.**
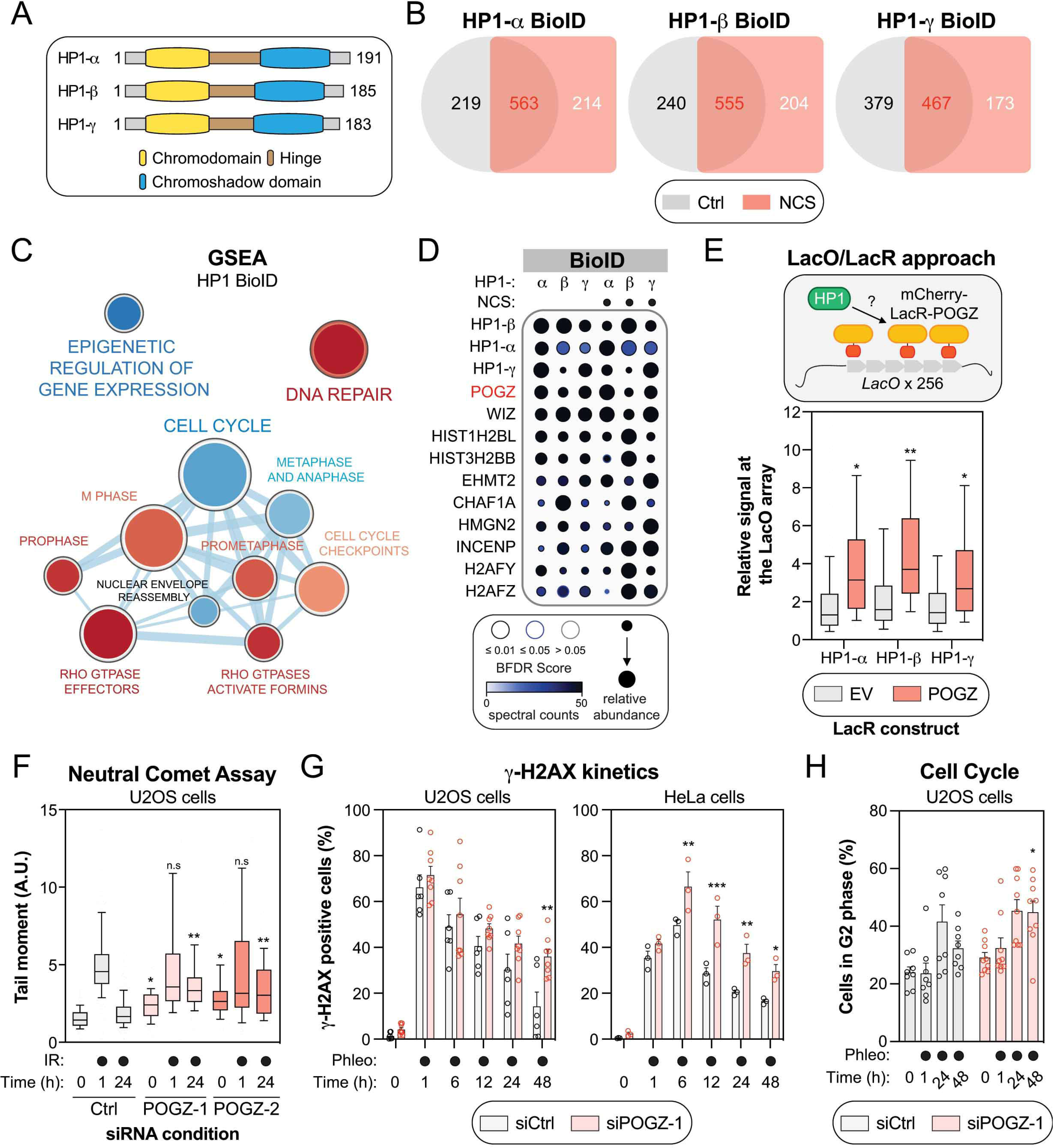
The HP1-interacting protein POGZ participates in DNA repair. (A) Schematic of the 3 human isoforms of the heterochromatin protein 1 (HP1). The N-terminal chromodomain of HP1 (in yellow) recognizes H3K9 methylation *in vitro*, with a preference for higher methylation states (H3K9me^1^>H3K9me^2^>H3K9me^3^), while the chromoshadow domain at its C-terminus (in blue) is involved in homo-/hetero-dimerization as well as interaction with other proteins containing a PXVXL motif. Both domains are separated by a hinge region (in brown). (B) Venn diagram outlining the distribution of HP1-α, -β and -γ high-confidence interactors identified by the BioID approach, under control (Ctrl; DMSO) or genotoxic conditions (NCS). (C) Gene set enrichment analysis (GSEA) visualization of the common HP1 high-confidence interactors using Reactome pathways. Enrichment maps were developed with a ranked interaction network (p < 0.2, FDR < 0.5 and overlap coefficient = 0.75) and the cell cycle cluster is provided in this panel. The complete interaction network can be found in Expanded View Fig.1D. Pathways enriched in: *(i)* control conditions are represented in blue; *(ii)* NCS-treated conditions are represented in red. (D) Dot plot of selected HP1 high-confidence interactors identified by BioID. The node size displays the relative abundance of a given prey across the 6 conditions analyzed, the node color represents the absolute spectral count sum (capped at 50 for display purposes), and the node edge color corresponds to the Bayesian False Discovery Rate (BFDR). Proteins were selected based on a SAINT score of > 0.95, BFDR of < 0.05, and ≥ 10 peptide count. (E) Schematic representing the LacO/LacR tethering system in U2OS cells (top panel). Box Quantification of the endogenous HP1 signal at the mCherry focus upon expression and tethering of mCherry-LacR alone (EV) or a construct containing POGZ. Data are represented as a box-and-whisker (10-90 percentile; bottom panel). At least 100 cells per condition were counted. Significance was determined by one-way ANOVA followed by a Tukey test. \**P<0.05*, \*\**P<0.0001*. (F) U2OS cells containing a non-targeting siRNA control (Ctrl), or one of two siRNA(s) against human POGZ (POGZ-1 or -2), were irradiated with 2 Gy before being collected at the indicated time points and assessed for comet tail migration in neutral conditions. Quantification of the neutral comet assay is represented as a box-and-whisker (10-90 percentile). At least 100 cells per condition were counted. Significance was determined by two-way ANOVA followed by a Dunnett’s test. \**P<0.05*, \*\**P<0.0001*. (G) U2OS (left panel, n=6) or HeLa cells (right panel, n = 3) were transfected with the indicated siRNA; 48 hours post-transfection, they were treated with the radiomimetic antibiotic, phleomycin (50 μg/mL), and collected at the indicated time points. Flow cytometry analysis of phosphorylated—H2AX signal was used to measure γ-H2AX endogenous signal. Data are represented as a bar graph showing the mean ± SEM, each replicate being represented by a round symbol. Significance was determined by two-way ANOVA followed by a Holm-Sidak’s test. \**P<0.05*, \*\**P<0.01*, \*\*\**P<0.0005*. (H) U2OS (n=8) were transfected with the indicated siRNA; 48 hours post-transfection, they were treated with the radiomimetic antibiotic, phleomycin (50 μg/mL), and collected at the indicated time points. Data are represented as a bar graph showing the mean ± SEM, each replicate being representing as a round symbol. Significance was determined by two-way ANOVA followed by a Sidak’s test. \**P<0.05*.

### POGZ modulates DNA repair *in vitro*

POGZ is a well-documented interactor of both HP1 and REV7/MAD2L2 (Vermeulen et al., 2010), and its contribution has been described during mitosis (Nozawa et al., 2010), but role to the maintenance of genome stability in interphase remains elusive (Baude et al., 2016). To confirm HP1-POGZ interaction in interphase, we took advantage of a single-cell assay where the association of two proteins can be assessed at an integrated LacO array by tethering a mCherry-LacR tagged version of a bait of interest (Fig.1E). This approach has been successful in recapitulating both BRCA1-PALB2 and RAD51D-XRCC2 interactions *in cellulo* (Orthwein, Noordermeer et al., 2015, Rivera, Di Iorio et al., 2017). Targeting a mCherry-LacR-tagged version of POGZ resulted in a significant accumulation of endogenous HP1-α, -β and –γ to the LacO array compared to control conditions (empty vector; EV; Fig.1E-F). To ascertain whether POGZ is relevant for DNA repair, we monitored the presence of DSBs in POGZ-depleted cells using the neutral comet assay (Lu, Liu et al., 2017). Interestingly, treatment of U2OS cells with two distinct small interfering RNAs (siRNAs) targeting POGZ (siPOGZ-1, and -2) resulted in a substantial increase of DSBs at steady-state (T0) compared to the control condition (siCtrl) (Fig.1F and Expanded View Fig.1G-H). Upon exposure to ionizing radiation (IR; 2 Gy), which generated a similar amount of DNA lesions in both siCtrl-and siPOGZ-treated cells (T=1hr), POGZ depletion caused a significant delay in the resolution of these DSBs at late time point (T=24hrs; Fig.1F), suggesting a role for POGZ during DNA repair. In parallel, we monitored the phosphorylation of the histone variant H2AX on serine 139 (γ-H2AX), which is a key step in the initiation of the response to DSBs and their subsequent repair. Using a well-established flow cytometry-based approach (Johansson, Fasth et al., 2017), POGZ depletion correlated with persistent phleomycin-induced DSBs (Phleo; 50 μg/mL) in both U2OS and HeLa cells compared to control conditions (siCtrl; Fig.1G and Expanded View Fig.1H). Importantly, we noted that this phenotype is associated with a prolonged G2 phase DNA damage checkpoint (Fig.1H), as well as the formation of micronuclei (Expanded View Fig. 1I-J), pointing towards an important role of POGZ in the maintenance of genome stability during genotoxic stress.

### POGZ promotes homology-based DNA repair pathways

To gain further insight into the contribution of POGZ *in vitro,* we generated both shRNA-mediated POGZ-depleted RPE1-hTERT cells (shPOGZ-1 and -2) and CRISPR-mediated POGZ-deleted HeLa cells (POGZΔ-1 and -2) (Expanded View Fig.2A). Based on our preliminary data, we hypothesized that depletion of POGZ should correlate with an increased cytotoxicity to DNA damaging agents that produce DSBs. Using the Sulforhodamine B (SRB) assay to determine cell density by measuring cellular protein content (Vichai & Kirtikara, 2006), we observed that POGZ-depleted cells were significantly more sensitive to the radiomimetic drug neocarzinostatin (NCS), irrespective of cell type (Fig.2A). Similar observations were made with the intercalating agent cisplatin (CIS) and the PARP inhibitor talazoparib (TZ) (Expanded View Fig.2B). Importantly, re-expression of a full-length mCherry-POGZ (FL) in POGZ-deleted HeLa cells (POGZΔ-1), ablated this hypersensitivity to both NCS and TZ (Fig.2B and Expanded View Fig.2C). Interestingly, dysregulated HR pathway has been linked to the cytotoxic potential of these drugs (Bhattacharyya, Ear et al., 2000, Farmer, McCabe et al., 2005, Yuan, Chang et al., 2002), suggesting a potential contribution of POGZ in this DNA repair pathway.

**Figure 2.**
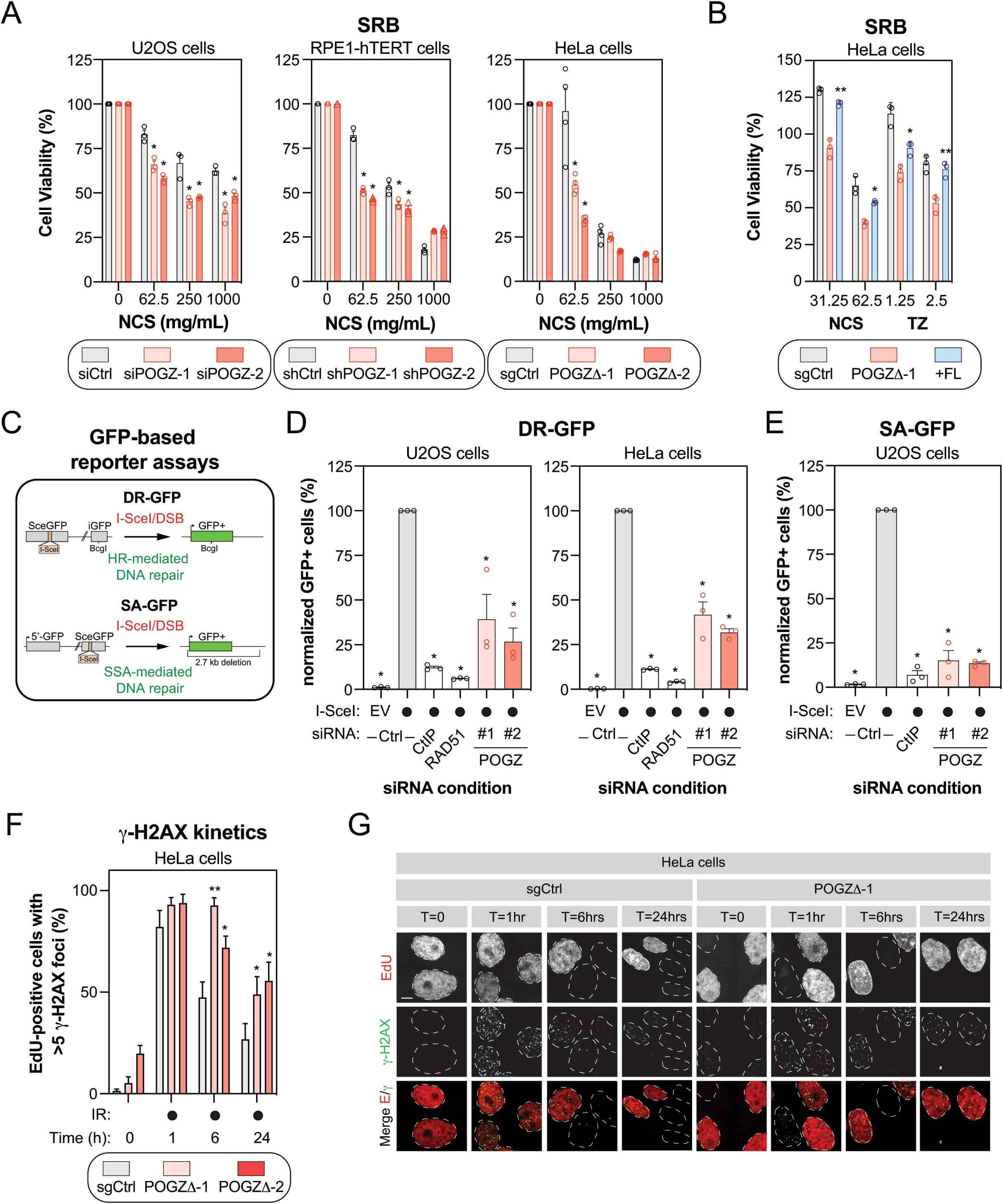
POGZ promotes homology-directed DNA repair pathways. (A) U2OS (left panel), RPE1-hTERT (middle panel), and HeLa cells (right panel) were monitored for their sensitivity to the radiomimetic drug NCS using the SRB assay. For each cell line, the following conditions were used: U2OS cells were transfected with a nontargeting siRNA (siCtrl) or an siRNA targeting human POGZ (siPOGZ-1 or -2); RPE1-hTERT cells were transduced a control shRNA (shCtrl) or a shRNA directed against human POGZ (shPOGZ-1 or -2); HeLa cells expressing a non-targeting sgRNA (sgCtrl) or a sgRNA targeting human POGZ and sub-cloned (POGZΔ-1 or -2). Cells were pulsed with NCS at the indicated concentrations for 1 hour, replenished with fresh medium and incubated for 4 days before being processed for SRB assays. Data are represented as a bar graph showing the relative mean ± SEM, each replicate being representing as a round symbol. Significance was determined by two-way ANOVA followed by a Bonferroni’s test. \**P<0.01*. (B) HeLa cells with (+FL, blue), or without full length POGZ-cDNA supplementation (POGZΔ-1, red), as well as control HeLa cells (sgCtrl, grey), were treated with NCS (1hr) or TZ (24h) at the indicated concentrations and processed as in (A) for SRB assay. Data are represented as a bar graph showing the relative mean ± SEM, each replicate being representing as a round symbol. Significance was determined by two-way ANOVA followed by a Bonferroni’s test. \**P<0.05*, \*\**P<0.005*. (C) Schematic diagram of the DR-GFP (top panel) and the SA-GFP (bottom panel) assays. (D) U2OS (left panel) and HeLa (right panel) cells containing the DR-GFP reporter construct were transfected with the indicated siRNA. 24h post-transfection. Cells were transfected with the I-SceI expression plasmid or an empty vector (EV), and the GFP+ population was analyzed 48h post-plasmid transfection. The percentage of GFP+ cells was determined for each individual condition and subsequently normalized to the non-targeting condition provided with I-SceI (siCtrl, I-SceI). Data are represented as the mean ± SEM, each replicate being representing as a round symbol (*n*=3). Significance was determined by one-way ANOVA followed by a Dunnett’s test. \**P≤0.0001*. (E) U2OS cells containing the SA-GFP reporter plasmid were processed and analyzed as in (D). Data are represented as the mean ± SEM, each replicate being representing as a round symbol (*n*=3). Significance was determined by one-way ANOVA followed by a Dunnett’s test. \**P≤0.0001*. (F) Quantification of γ-H2AX foci in HeLa cells where POGZ has been targeted by CRISPR technology (POGZΔ-1 or -2) and in control HeLa cells (sgCtrl). Cells were exposed to 1 Gy before being pulsed with Edu for 1hr and were recovered at the indicated time points. Cells were fixed, stained, and imaged via confocal microscopy. Data are the percent of EdU+ cells in a field of view with >5 γ-H2AX foci and represented as a bar graph showing the mean ± SEM (*n* = 3, with at least 200 cells analyzed for each time point). Significance was determined by two-way ANOVA followed by a Dunnett’s test. \**P<0.05,* \*\**P<0.0005*. (G) Representative images used for quantification in (F). Scale bar = 5 μm.

To directly test this possibility, we employed a well-established GFP-based reporter assay that monitors DNA repair by HR, the DR-GFP assay, and evaluated the impact of POGZ depletion on restoring GFP expression (Fig.2C). As positive controls, we introduced siRNAs targeting key components of the HR pathway: CtIP and RAD51. In both U2OS and HeLa DR-GFP cells, POGZ depletion led to a significant decrease in HR using two distinct siRNAs (Fig.2D). Next, we evaluated whether POGZ could regulate another homology-based DNA repair pathway, SSA, using the SA-GFP assay (Fig.2C). We observed a similar phenotype in the SA-GFP reporter assay, where POGZ depletion resulted in a significant reduction of the GFP signal (Fig.2E), indicative of a reduced SSA efficiency. These results are consistent with a model where POGZ promotes homology-directed DNA repair pathways.

If our model is correct, loss of POGZ should impair the resolution of DSBs in S/G2 phases of the cell cycle, where HR is restricted. We therefore pulse-labelled our cell lines with 5-ethynyl-2’-deoxyuridine (EdU) and monitored the presence of IR-induced γ-H2AX foci in EdU-positive (EdU+) cells by immunofluorescence. As predicted, the proportion of γ-H2AX+/EdU+ cells (>5 γ-H2AX foci) remained significantly higher upon partial (U2OS and RPE1-hTERT cells) or complete loss of POGZ (HeLa cells) at late time points (6 and 24hrs) compared to control conditions (Fig.2F-G, Expanded View Fig.2D). As previously observed, this phenotype correlated with a sustained G2-phase cell cycle checkpoint in POGZ-depleted cells upon exposure to IR (Expanded View Fig.2E), likely due to the sustained phosphorylation of ATM (Ser1981) and CHK1 (Ser345) following exposure to DNA damage (Expanded View Fig.3A-B). Interestingly, the lack of a sustained activation of ATR (pATR Ser1989) would suggest that POGZ promotes HR prior to the formation of ssDNA during DNA end resection (Expanded View Fig.3A). Of note, upon exposure to IR, we noted a significantly larger proportion of apoptotic cells in POGZ-depleted versus control conditions, as monitored by the Annexin V binding (Expanded View Fig.3C). Our data indicate that POGZ plays a pivotal role in regulating both DNA repair by HR and the associated G2 DNA damage checkpoint, thereby modulating permanent cell fate such as apoptosis.

### POGZ facilitates the accumulation of BRCA1 at DSBs

To pinpoint at which step(s) POGZ is regulating HR-mediated DNA repair, we monitored the impact of its depletion on the recruitment of well-established DSB signaling factors. Here, we used the LacO-LacR system where we expressed a mCherry-LacR-tagged version of the endonuclease FokI (Expanded View Fig.3D), allowing the formation of localized DSBs and the visualization of DNA repair factors at the LacO repeat by immunofluorescence (Shanbhag & Greenberg, 2013). As expected, we did not observe any significant impact on the recruitment of 53BP1 and RIF1 to FokI-induced DSBs upon depletion of POGZ (siPOGZ-1 and -2; Fig.3A and Expanded View Fig.3E). However, under the same conditions, the presence of BRCA1 and RAD51 at DSBs was significantly impaired (Fig.3A and Expanded View Fig.3E). In fact, we made similar observations when we monitored the formation of IR-induced BRCA1 foci in cells pulsed with EdU (Fig.3B-C), pointing towards a key role of POGZ in regulating BRCA1 accumulation at DNA damage sites.

**Figure 3.**
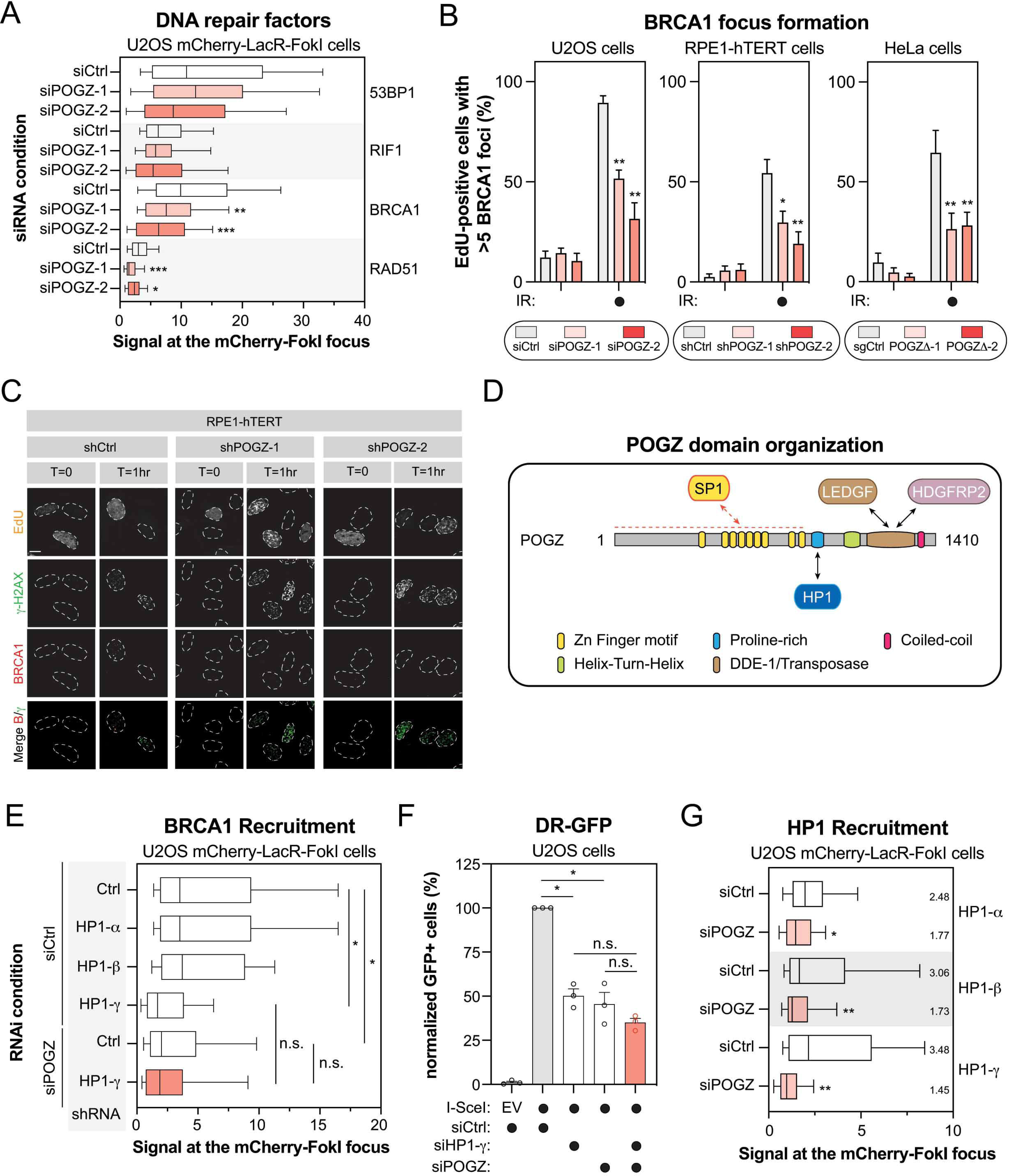
POGZ modulates the presence of BRCA1 at DNA damage sites. (A) U2OS stably expressing mCherry-LacR-FokI were transfected with the indicated siRNA. 24h post-transfection, DNA damage was induced using Shield1 and 4-OHT. Immunofluorescence against the indicated DNA repair proteins was subsequently performed to monitor their accumulation at sites of DNA damage by confocal microscopy. Data are represented as a box-and-whisker (10-90 percentile) where the fluorescent signal at the mCherry focus was normalized to nuclear background. At least 100 cells per condition were counted (n=3). Significance was determined by two-way ANOVA followed by a Dunnett’s test. \**P<0.05*, \*\**P<0.005,* \*\**P≤0.0005*. (B) U2OS (left panel), RPE1-hTERT (middle panel), and HeLa cells (right panel) were monitored for their capacity to form IR-induced BRCA1 foci. For each cell line, the following conditions were used: U2OS cells were transfected with a nontargeting siRNA (siCtrl) or an siRNA targeting human POGZ (siPOGZ-1 or -2); RPE1-hTERT cells were transduced a control shRNA (shCtrl) or a shRNA directed against human POGZ (shPOGZ-1 or -2); HeLa cells were expressing a non-targeting sgRNA (sgCtrl) or a sgRNA targeting human POGZ and sub-cloned (POGZΔ-1 or -2). Cells were exposed to 1 Gy before being pulsed with Edu for 1hr and were recovered 1h post-exposure to IR. Cells were fixed, stained, and imaged via confocal microscopy. Data are the percent of EdU+ cells in a field of view with >5 BRCA1 foci and represented as a bar graph showing the mean ± SEM (*n* = 3, with at least 100 cells analyzed for each time point). Significance was determined by two-way ANOVA followed by a Dunnett’s test. \**P<0.005,* \*\**P<0.001*. (C) Representative images used for quantification in (B). Scale bar = 5 μm. (D) Schematic diagram outlining the different domains of human POGZ. Each structural domain and interacting partners are indicated. (E) U2OS stably expressing mCherry-LacR-FokI were transduced with the indicated shRNA and subsequently transfected with the indicated siRNA. 24h post-transfection, DNA damage was induced using Shield-1 and 4-OHT. Immunofluorescence against BRCA1 was subsequently performed to monitor its accumulation at sites of DNA damage by confocal microscopy. Data are represented as a box-and-whisker (10-90 percentile) where the fluorescent signal at the mCherry focus was normalized to nuclear background. At least 50 cells per condition were counted. Significance was determined by two-way ANOVA followed by a Dunnett’s test. \**P<0.0001*. (F) U2OS cells containing the DR-GFP reporter construct were transfected with the indicated siRNA. 24h post-transfection. Cells were transfected with the I-SceI expression plasmid or an empty vector (EV), and the GFP+ population was analyzed 48h post-plasmid transfection. The percentage of GFP+ cells was determined for each individual condition and subsequently normalized to the non-targeting condition provided with I-SceI (siCtrl, I-SceI). Data are represented as a bar graph showing the relative mean ± SEM, each replicate being representing as a round symbol (n=3). Significance was determined by one-way ANOVA followed by a Dunnett’s test. \**P≤0.0001*. (G) U2OS stably expressing mCherry-LacR-FokI were transduced with the indicated siRNA. 24h post-transfection, DNA damage was induced using Shield-1 and 4-OHT. Immunofluorescence against the indicated HP1 isoform was subsequently performed to monitor its respective accumulation at sites of DNA damage by confocal microscopy. Data are represented as a box-and-whisker (10-90 percentile) where the fluorescent signal at the mCherry focus was normalized to nuclear background. At least 75 cells per condition were counted. Significance was determined by two-way ANOVA followed by a Dunnett’s test. \**P<0.005,* \*\**P<0.0001*.

POGZ is a 1410 amino-acid protein with multiple functional domains (Fig.3D): one atypical and 8 classical C_2_H_2_ zinc finger domains at its N-terminus, a proline-rich domain, a helix-turn-helix domain also identified as a centromere protein (CENP) B-like DNA binding domain, a putative DDE-1 transposase domain and a coiled-coil motif at its C-terminus. POGZ has been proposed to act as a negative regulator of gene expression in different biological processes, including hematopoiesis and neuronal development (Gudmundsdottir, Gudmundsson et al., 2018, Matsumura, Seiriki et al., 2020, Suliman-Lavie, Title et al., 2020). To exclude the possibility that POGZ may indirectly control DNA repair through its known transcriptional function, we profiled by quantitative RT-PCR (qPCR) the expression of BRCA1 and BARD1 as well as downstream effectors in the HR pathway, including PALB2, BRCA2 and RAD51. However, we did not notice any substantial changes in their expression upon depletion of POGZ in both HeLa and RPE1-hTERT cells (Expanded View Fig.4A). We therefore explored the possibility that POGZ may have a more direct contribution to DNA repair by monitoring its recruitment to DNA damage sites. Interestingly, we observed that HA-tagged POGZ rapidly accumulates at laser micro-irradiation-induced DNA damage in U2OS cells, co-localizing with γ-H2AX (Expanded View Fig.4B-C). This recruitment is a transient event and POGZ accumulation disappears within 30 min after laser micro-irradiation, a dynamic reminiscent of what has been observed with HP1 (Ayoub et al., 2008, Baldeyron et al., 2011, Luijsterburg et al., 2009, Zarebski et al., 2009).

### POGZ allows the presence of the HP1-γ/BARD1/BRCA1 complex at DSBs

HP1 is a well-established factor in the response to DSBs (Bartova, Malyskova et al., 2017), at least, in part through the retention of BRCA1 at DNA damage sites by HP1-γ (Wu et al., 2015). We confirmed these observations in the U2OS mCherry-LacR-FokI cells where depletion of HP1-γ, but neither HP1-α or -β significantly impaired the accumulation of BRCA1 at the focal DNA damage site (Fig.3E and Expanded View Fig.4D-E). If POGZ facilitates BRCA1 retention at DSBs in a HP1-γ-dependent manner, co-depletion of both POGZ and HP1-γ should not impair further BRCA1 accumulation at DSBs any further. Indeed, we did not notice any significant decrease in the retention of BRCA1 in this condition compared to either HP1-γ or POGZ depletion alone (Fig.3E and Expanded View Fig.4D-E). Similar observations were made in the DR-GFP assay where co-depletion of both POGZ and HP1–γ did not further decrease HR-mediated DNA repair compared to the individual HP1–γ or POGZ depletion in the U2OS cell line (Fig.3F). Interestingly, POGZ depletion significantly impaired the recruitment of all HP1 isoforms to FokI-induced DSBs; however, we noted that it had the most drastic impact on HP1-γ (Fig.3G and Expanded View Fig.4F), pointing towards a model where POGZ promotes HR by allowing the accumulation of BRCA1 at DSBs in an HP1-γ-dependent manner.

To gain further insight into how POGZ promotes DNA repair, we performed a structure-function analysis by generating several truncation mutants (Fig.4A). First, we tested their sub-cellular localization by expressing a mCherry-tagged version of these mutants in HEK293T cells. Mutants lacking the proline-rich domain (amino acids 800-848; ΔHP1), which mediates HP1 binding (Nozawa et al., 2010), were unable to accumulate in the nucleus (Expanded View Fig.4G-H), confirming a previous report linking the nuclear accumulation of POGZ to its ability to bind to HP1(Matsumura et al., 2020). We first focused our attention on the POGZ^366-848^ mutant, which contains the zinc finger region of POGZ as well as its HP1 binding domain and retains the capacity to localize to the nucleus (Expanded View Fig.4G-H). We re-expressed a Flag-tagged version in this mutant in POGZ-depleted U2OS cells containing the inducible LacR-FokI system and monitored its ability to restore BRCA1 recruitment to DSBs. Strikingly, POGZ^366-848^ mutant restored BRCA1 accumulation to localized DSBs, at a level comparable to full-length (FL) POGZ (Fig.4B and Expanded View Fig.5A). Interestingly, this mutant allowed the recruitment of both BARD1 and HP1-γ to the localized DNA damage site (Fig.4B and Expanded View Fig.5A). To further dissect the critical domain(s) of POGZ promoting DNA repair, we generated a construct exclusively containing the HP1 binding domain (HPZ) and expressed it in HeLa POGZΔ cells. We pulse-labelled these cells with EdU and monitored their capacity to form IR-induced BRCA1 foci by immunofluorescence. Remarkably, the POGZ HPZ mutant restored BRCA1 foci in POGZ-deleted HeLa cells, to a similar extent as FL POGZ (FL; Fig.4C). Critically, we observed that HPZ-expressing cells display significantly less IR-induced γ-H2AX foci over time, compared to HeLa POGZΔ cells (Fig.4E-F and Expanded View Fig.5B), suggesting that this construct is able to restore DSB resolution. In light of these data, we conclude that the HP1 binding domain of POGZ is necessary and sufficient to promote BRCA1 recruitment to DSBs and HR-mediated DNA repair, primarily through HP1-γ.

**Figure 4.**
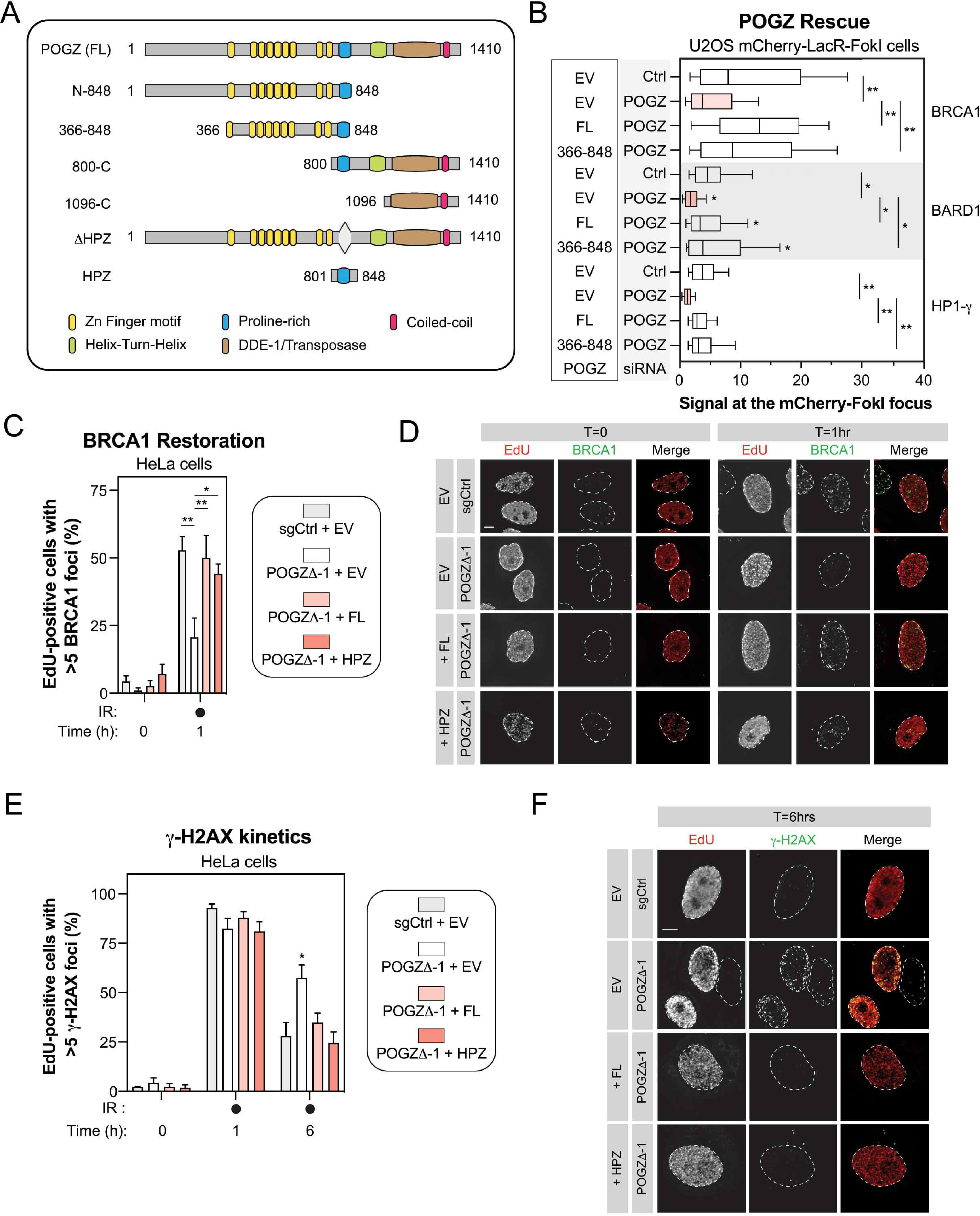
POGZ mediates BRCA1/BARD1 accumulation at DSBs through its HP1-binding site. (A) Schematic diagram outlining the different domains of human POGZ and the different deletion constructs of POGZ that we generated and analyzed. (B) U2OS stably expressing mCherry-LacR-FokI were transduced with the indicated siRNA. 24h post-transfection, cells were transfected with an empty Flag vector (EV) or a siRNA-resistant FLAG-tagged POGZ cDNA construct corresponding to indicated rescue mutant (full-length, FL; POGZ^366-848^, 366-848). 24h after plasmid transfection, DNA damage was induced using Shield-1 and 4-OHT. Immunofluorescence against the indicated protein was subsequently performed to monitor its respective accumulation at sites of DNA damage by confocal microscopy. Data are represented as a box-and-whisker (10-90 percentile) where the fluorescent signal at the mCherry focus was normalized to nuclear background. At least 100 cells per condition were counted. Significance was determined by two-way ANOVA followed by a Dunnett’s test. \**P<0.005,* \*\**P<0.0001*. (C) HeLa cells transfected with the indicated construct were monitored for their capacity to form IR-induced BRCA1 foci. HeLa cells where POGZ has been targeted by CRISPR technology (POGZΔ-1) and in control HeLa cells (sgCtrl) were transfected by an empty Flag vector (EV) or a sgRNA-resistant FLAG-tagged POGZ cDNA construct corresponding to indicated rescue mutant (full-length, FL; POGZ^801-848^, HPZ). Cells were exposed to 1 Gy before being pulsed with Edu for 1hr and were recovered 1h post-exposure to IR. Cells were fixed, stained, and imaged via confocal microscopy. Data are the percent of EdU+ cells in a field of view with >5 BRCA1 foci and represented as a bar graph showing the mean ± SEM (*n* = 3, with at least 200 cells analyzed for each time point). Significance was determined by two-way ANOVA followed by a Dunnett’s test. \**P<0.05,* \*\**P<0.0001*. (D) Representative images used for quantification in (C). Scale bar = 5 μm. (E) Similar as in (C) except that γ-H2AX foci were monitored at the indicated time points by confocal microscopy. Data are the percent of EdU+ cells in a field of view with >5 γ-H2AX foci and represented as a bar graph showing the mean ± SEM (*n* = 3, with at least 200 cells analyzed for each time point). Significance was determined by two-way ANOVA followed by a Dunnett’s test. \**P<0.05*. (F) Representative images used for quantification in (E). Scale bar = 5 μm.

### Loss of *Pogz* impairs proper murine development *in vivo*

Heterozygous pathogenic variants in *POGZ* have been linked to a rare human disorder, known as the White-Sutton syndrome (WHSUS), characterized by craniofacial abnormalities such as microcephaly, a developmental delay, intellectual disabilities as well as behavioural problems (e.g. hyperactivity, overly friendly behaviour), and in certain rare cases, recurrent infections (Dentici et al., 2017, Ferretti et al., 2019, Fukai et al., 2015, Matsumura et al., 2016, Samanta et al., 2020, Tan et al., 2016, Ye et al., 2015, Zhao et al., 2019a, Zhao et al., 2019b). These observations prompted us to define the relevance of POGZ *in vivo*.

Using the CRISPR/Cas9 technology, we targeted exons 9 and 10 of *Pogz* in embryonic stem (ES) cells derived from C57BL/6J mice in order to generate a *Pogz* knock-out mouse model (Fig.5A). Remarkably, we failed to obtain any viable constitutive *Pogz* knockout mice (*Pogz*^Δ/Δ^) (Fig.5B). Most homozygous *Pogz* knock-out embryos died at mid-to late gestation, between embryonic day E12.5 and E14.5 (Fig.5C), suggesting that *Pogz* is essential for mouse embryonic development. However, we were successful in deriving mouse embryonic fibroblasts (MEFs) from E12.5 embryos where we confirmed that CRISPR-mediated targeting of one or both alleles of *Pogz* resulted in its partial or complete ablation at the protein level, respectively (Fig.5D). As heterozygous pathogenic mutations in *POGZ* lead to the WHSUS, we decided to investigate whether *Pogz* haplo-insufficiency (*Pogz*^+/Δ^) may have an overall impact on murine development. Indeed, *Pogz^+/Δ^* mice have a significantly lower body weight compared to their wild-type counterparts as early as 4 weeks post-birth (Fig.5E). This growth defect correlated with smaller organs in *Pogz^+/Δ^* mice, including the thymus, the spleen, the liver and the brain (Fig.5F), suggesting that Pogz levels need to be tightly regulated to allow proper murine development.

**Figure 5.**
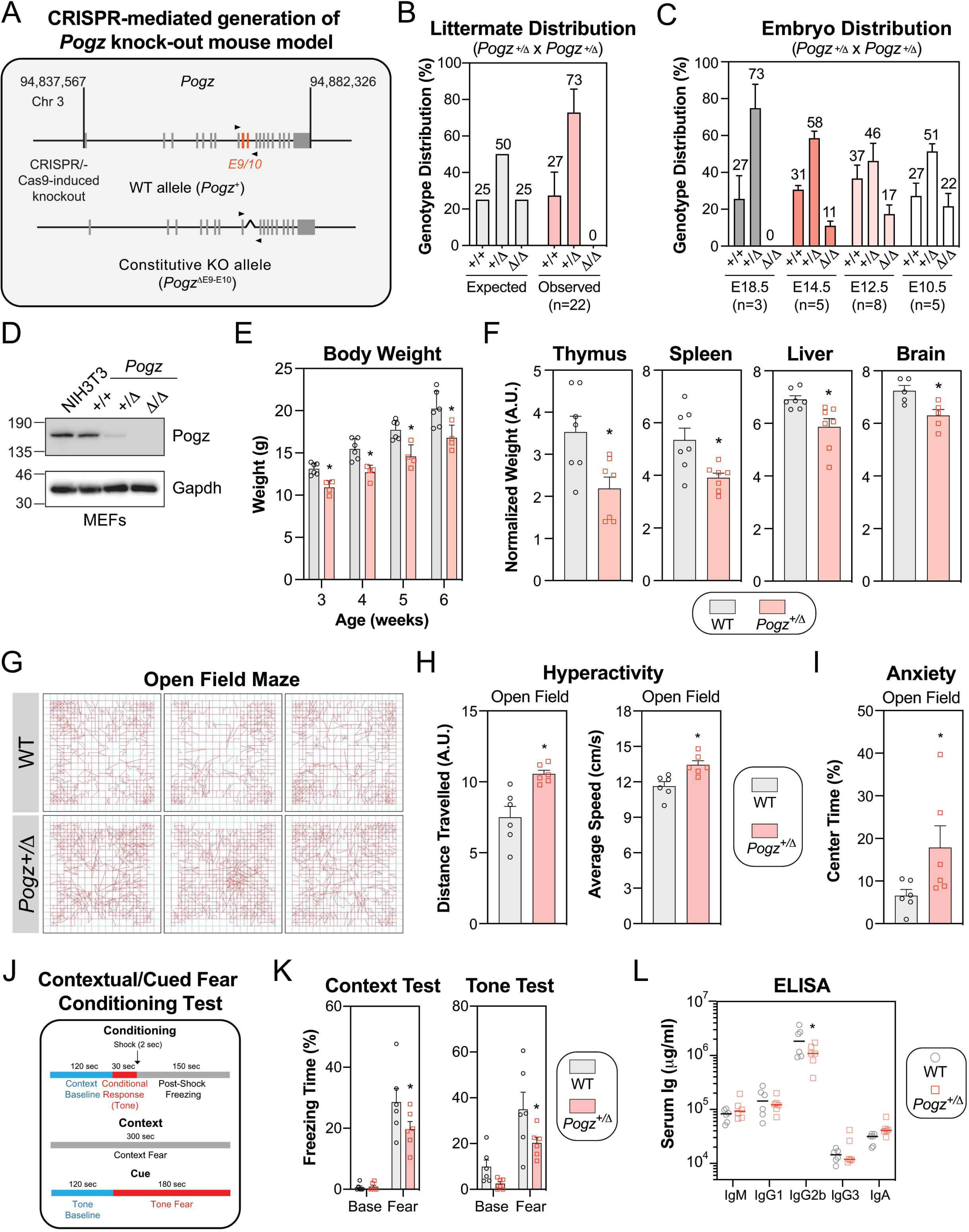
Pogz haplo-insufficiency in mice recapitulates the clinical features observed in patients affected by the WHSUS. (A) Schematic diagram outlining the generation of CRISPR/Cas9-mediated *Pogz^+/Δ^* mouse model. A region spanning critical exons 9 and 10 of the murine *Pogz* gene on chromosome 3 was deleted using dual sgRNA CRISPR. (B) Expected and observed genotypic distribution of offspring of heterozygous *Pogz^+/Δ^* crosses. Genotype was determined by PCR at time of weaning (3 weeks). Data are represented as a bar graph showing the mean ± SEM (n=22 individual litters, across 5 different breeding pairs). (C) Observed genotypic distribution of offspring of heterozygous *Pogz^+/Δ^* crosses at specified embryonic day. Each litter is considered a biological replicate. Data are represented as a bar graph showing the mean ± SEM (n for each embryonic day is specified). (D) Expression analysis of Pogz by western blot in mouse embryonic fibroblasts (MEFs) generated from E12.5 embryos with the indicated genotype. NIH3T3 cells were used as a comparison. Gadph was used a loading control. (E) The body mass of male wild-type (WT) or *Pogz^+/Δ^* mice was monitored weekly for 4 weeks post-weaning. Data are represented as a bar graph showing the mean ± SEM and each mouse is represented by a round dot. At least 4 mice per genotype was monitored. Significance was determined by two-way ANOVA followed by a Sidak’s test. \**P<0.05*. (F) The indicated organ mass of male wild-type (WT) or *Pogz^+/Δ^* mice was calculated relative to total body mass. Data are represented as a bar graph showing the mean ± SEM and each mouse is represented by a round dot (n=7). Significance was determined by unpaired two-tailed t-test. \**P<0.05*. (G) Representative movement traces of the indicated mice used for quantification in (H) and (I). (H) Quantification of the distance travelled (left panel) and the average speed (right panel) of each mouse (n=6 for each genotype) in the open field maze. Data are represented as a bar graph showing the mean ± SEM and each mouse is represented by a round dot. Significance was determined by unpaired two-tailed t-test. \**P<0.005*. (I) The percentage of time that each mouse spent in the middle of the open field was quantified and represented as the mean ± SEM, each mouse being represented by a round dot (n = 6 mice per genotype). Significance was determined by unpaired two-tailed t-test. \**P<0.005*. (J) Schematic diagram outlining the conditioning/experimental set up quantified in (K) of the contextual fear tests. (K) The percentage of freezing time in the different experimental conditions (context test, left panel; tone test, right panel) was monitored for each mouse and is represented as the mean ± SEM, each mouse being represented by a round dot (n=6 mice per genotype). Significance was determined by two-way ANOVA followed by a Sidak’s test. \**P<0.05*. (L) Plasma was isolated from cardiac punctures of wild-type (WT) or *Pogz^+/Δ^* mice (8 weeks) and assessed for circulating levels of specified immunoglobulin isotypes (n=6 mice per genotype). Significance was determined by two-way ANOVA followed by a Bonferroni’s test. \**P<0.005*.

As microcephaly often correlates with impaired cognitive functions (Pavone, Pratico et al., 2017), we performed a systematic behavioral analysis of our mouse model. Using the open field maze test, we noticed that *Pogz^+/Δ^* mice have significantly more locomotor activity, with a substantially greater average speed, than their age-matched wild-type littermates (Fig.5G-H), indicative of a hyperactive behavior. Furthermore, *Pogz* haplo-insufficient mice spent more time in the center of the arena (inner zone of the maze), which is typically linked to a decreased level of anxiety (Fig. 5G-I). To assess the intellectual capabilities of our *Pogz^+/Δ^* mice, we performed a contextual and cued fear conditioning test (Fig.5J). In this procedure, a neutral conditioned stimulus (i.e. steady tone) is paired with an aversive unconditioned stimulus (mild foot shock) and the time during which animals present a lack of mobility or reduced locomotor activity is a readout of learning/memory performances. Interestingly, *Pogz* haplo-insufficient mice exhibited decreased freezing behavior compared to control mice, in which the baseline behavior remains comparable (Fig.5K), indicative of a deficit in contextual learning and memory capacities.

Severe cases of the WHSUS are characterized by recurrent infections (Stessman, Willemsen et al., 2016), likely due to a dysfunctional immune system. For instance, reduced lymphoid organ weight, as observed in *Pogz^+/Δ^* mice (Thymus, Spleen; Fig.5F), may be indicative of a compromised immune system. We evaluated whether *Pogz* haplo-insufficient mice have a compromised humoral immune response by measuring total immunoglobin (Ig) levels of IgM, IgG1, IgG2b, IgG3 and IgA in the serum. Interestingly, we found that serum IgG2b levels were almost 50% lower in *Pogz^+/Δ^* mice compared to age-matched wild-type controls (Fig.5L), indicative of a defective antibody response. Importantly, patients with a selective IgG2 subclass deficiency usually suffer from recurrent upper and lower respiratory tract infections (Barton, Barton et al., 2020), which have also been reported in severe cases of WHSUS (Stessman et al., 2016). Altogether, our data indicate that *Pogz* haplo-insufficient mice present a developmental delay, a deficit in intellectual abilities, a hyperactive behaviour as well as a compromised humoral immune response, reminiscent of the clinical features reported in WHSUS patients.

### *Pogz* haplo-insufficient mice display features of genomic instability

In light of our findings *in vitro*, we wondered whether *Pogz^+/Δ^* mice present any DNA repair defect. To test this hypothesis, we exposed mice to IR (8 Gy) and monitored their survival over time. Interestingly, IR-induced mortality occurred significantly faster in *Pogz^+/Δ^* mice than in wild-type controls (median survival: 7 days vs. 11 days; Fig.6A), suggesting that Pogz may also be relevant to promote DNA repair *in vivo*. We therefore tested whether *Pogz^+/Δ^* mice spontaneously accumulate DSBs by monitoring both γ-H2AX levels and intensity in the spleen by flow cytometry. Strikingly, we observed significantly more γ-H2AX-positive splenocytes in *Pogz^+/Δ^* mice compared to controls (WT; Fig.6B and Expanded View Fig.6A), and they display substantiality more γ-H2AX signal intensity than wild-type (Fig.6B and Expanded View Fig.6A). We extended our analysis to the brain, where we stained sections of the cerebral cortex for γ-H2AX (Fig.6C and Expanded View Fig.6B). Again, we noted substantially more γ-H2AX-positive cells in *Pogz^+/Δ^* mice vs. wild-type in this region of the brain (Fig.6C and Expanded View Fig.6B), pointing towards a global DNA repair defect *in vivo*.

**Figure 6.**
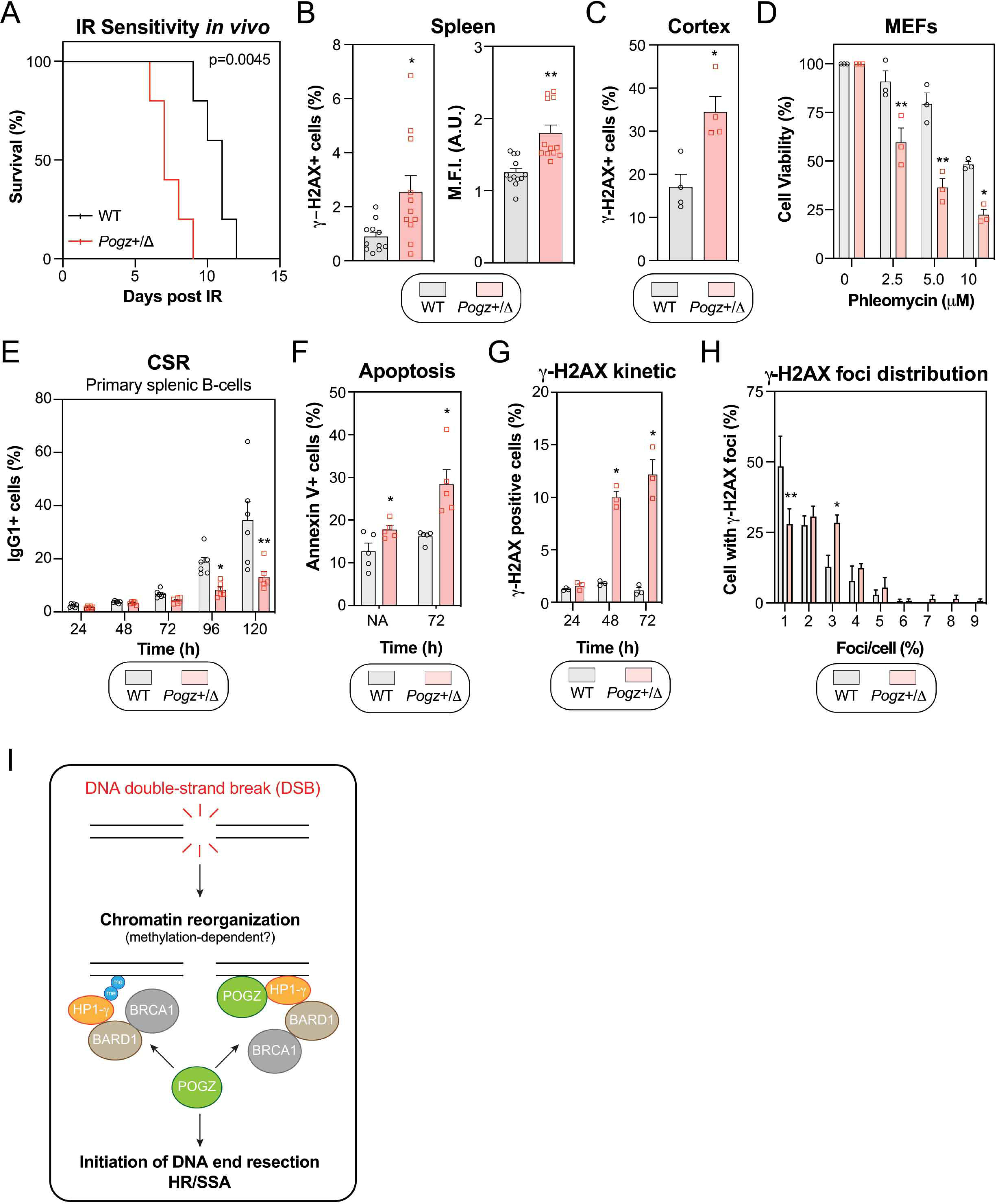
Pogz haploinsufficiency correlates with features of impaired DNA repair in mice. (A) Wild-type (WT) and *Pogz^+/Δ^* mice were subjected to a lethal dose of ionizing radiation (8.5 Gy) before recovering in sterile conditions and being assessed for their sensitivity to IR. Data are represented as a Kaplan-Meier survival curve of each genotype ((n=6 mice per genotype). Significance was determined by log-rank (Mantel-Cox) test. \**P<0.005*. (B) Quantification of phosphorylated-H2AX (γ-H2AX) levels by flow cytometry. Splenocytes isolated from 8-week-old wild-type (WT) and *Pogz^+/Δ^* mice were processed for γ-H2AX staining and data are represented as bar graph showing the mean percentage of cells that were γ-H2AX-positive ± SEM (left panel) or the mean fluorescence intensity (M.F.I.) of the γ-H2AX signal ± SEM (right panel), each mouse being represented by a round dot (n = 11 for each genotype). Significance was determined by unpaired two-tailed t-test. \**P<0.05,* \*\**P<0.0005*. (C) Brain slices were sectioned from 6-week-old wild-type (WT) and *Pogz^+/Δ^* mice, followed by immunostaining for phosphorylated H2AX (γ-H2AX). Data are the percentage of cells with γ-H2AX signal present in the nucleus per field of view and are represented as a bar graph showing the mean ± SEM (n= 4 for each genotype with three distinct fields quantified for each mouse). (D) MEFs were monitored for their sensitivity to the radiomimetic drug phleomycin using the SRB assay. Immortalized MEFS obtained from the indicated genotype were treated with increasing concentrations of phleomycin for 1hr, replenished with fresh medium and incubated for 4 days before being processed for SRB assays. Data are represented as a bar graph showing the relative mean ± SEM, each replicate being representing as a round symbol. Significance was determined by two-way ANOVA followed by a Bonferroni’s test. \**P<0.005*, \*\**P<0.0005*. (E) CD43-negative primary splenocytes from 8-week-old wild-type (WT) and *Pogz^+/Δ^* mice were stimulated *ex vivo* with IL-4 (50 ng/mL) and LPS (25 μg/mL). Cells were harvested at the indicated time points and assessed for their surface expression of IgG1 by flow cytometry. Data are represented as a bar graph showing the mean ± SEM, each mouse being represented by a round (WT) or square (*Pogz^+/Δ^)* dot (n=6 for each genotype). Significance was determined by two-way ANOVA followed by a Bonferroni’s test. \**P<0.05*, \*\**P<0.0001*. (F) Similar as in (E), except that cells were monitored for apoptosis by Annexin V staining. Data are represented as a bar graph showing the mean ± SEM, each mouse being represented by a round (WT) or square (*Pogz^+/Δ^)* dot (n=5 for each genotype). Significance was determined by two-way ANOVA followed by a Bonferroni’s test. \**P<0.05*. (G) Similar as in (E), except that cells were monitored for phosphorylated H2AX (γ-H2AX) levels by flow cytometry at the indicate time points post-stimulation with IL-4/LPS. Data are represented as a bar graph showing the mean ± SEM, each mouse being represented by a round (WT) or square (*Pogz^+/Δ^)* dot (n=3 for each genotype). Significance was determined by two-way ANOVA followed by a Bonferroni’s test. \**P<0.05*. (H) Stimulated CD43-negative splenocytes were loaded on slides via Cytospin and processed for γ-H2AX immunofluorescence. Cells were fixed, stained, and imaged via confocal microscopy. Data are the percentage of cells in a field of view with indicated γ-H2AX foci and are represented as a bar graph showing the mean ± SEM. At least 100 cells per genotype were counted. Significance was determined by two-way ANOVA followed by a Sidak’s test. \**P<0.05,* \*\**P<0.005*. (I) Schematic diagram outlining how POGZ is regulating the repair of DSBs by HR.

To validate our observations *in vivo*, we took advantage of MEFs that we derived from our mouse model (Fig.5D) and we tested their sensitivity to two distinct DNA damaging drugs, phleomycin and camptothecin. As expected, *Pogz^+/Δ^* MEFs were hypersensitive to both drugs compared to wild-type in the SRB assay (Fig.6D and Expanded View Fig.6C), confirming that partial loss of Pogz impairs DNA repair. *Ex vivo* stimulation of B-cells is another well-established system where the integrity of DNA repair pathways can be examined. Importantly, intact DNA repair pathways are required for the resolution of activation-induced deaminase (AID)-induced DSBs during class switch recombination (CSR; Expanded View Fig.6D). We therefore extracted primary splenic B-cells from both WT and *Pogz^+/Δ^* mice and activated them *ex vivo* with a cocktail of cytokines (IL-4/LPS) to induce class switching from IgM to IgG1. Interestingly, we noted that partial loss of *Pogz* significantly impaired CSR at both 96h and 120h post-activation (Fig.6E and Expanded View Fig.6E). Importantly, we confirmed that this phenotype was neither due to a proliferation defect (Expanded View Fig.6F), nor a lack of *Aicda* expression (Expanded View Fig.6G). As expected, we observed a significant reduction (∼50%) of *Pogz* mRNA expression in *Pogz^+/Δ^* mice compared to wild-type (Expanded View Fig.6G). Successful CSR relies on the capacity to resolve on-target DSBs by NHEJ and alternative end-joining (a-EJ) pathways in G1 phase of the cell cycle (Expanded View Fig.6D) (Cortizas, Zahn et al., 2013). However, a functional HR pathway has been shown to be important in the repair of both on-and off-target DSBs, thereby promoting B-cell survival (Expanded View Fig.6D) (Hasham, Donghia et al., 2010). Based on our data *in vitro*, we wondered whether *Pogz^+/Δ^* B-cells attempting CSR might be more prone to apoptosis, explaining the apparent deficit of CSR. In fact, we observed a substantial increase in the proportion of Annexin V-positive *Pogz^+/Δ^* B-cells compared to control conditions upon activation with a cocktail of cytokines (Fig.6F and Expanded View Fig.6H). Of note, a significant fraction of non-activated (NA) *Pogz^+/Δ^* B-cells were Annexin V-positive (Fig.6F), likely linked to the γ-H2AX signal that we observed previously in splenocytes (Fig.6B and Expanded View Fig.6A). If our model is right, *Pogz^+/Δ^* B-cells should accumulate a significant amount of DSBs upon induction of CSR. We therefore monitored γ-H2AX levels by flow cytometry and we noted that a significant increase in the proportion of γ-H2AX-positive *Pogz^+/Δ^* B-cells compared to wild-type at 48h and 72h post-activation (Fig.6G). We also monitored the number of γ-H2AX foci/cell in both wild-type and *Pogz^+/Δ^* B-cells upon induction of CSR. As expected, B-cell activation resulted in a significant increase in the proportion of cells displaying γ-H2AX foci (Expanded View Fig.6I). Interestingly, around half of these activated B-cells contain more than 2 γ-H2AX foci in a haplo-insufficient background compared to ∼25% in a wild-type context (Fig. 6H and Expanded View Fig.6J-K), suggesting that Pogz is critical for the repair of off-target DSBs during CSR. Altogether, our data are consistent with a model where POGZ promotes the repair of DSBs by homology-directed DNA repair pathways both *in vitro* and *in vivo* (Fig.6I).

## DISCUSSION

DNA repair by HR is an essential process for both the maintenance of genome stability and the generation of genetic diversity. A major factor influencing the commitment of a DSB to HR relates to its cell cycle positioning and the availability of a donor template. However, several additional elements participate in this “decision-making” process, including the presence of the BRCA1/BARD1 complex at DSBs, which is influenced by the epigenetic context. For instance, the direct recognition of H4K20me^0^ by the ankyrin repeat domain of BARD1 allows BRCA1 to be present on newly synthesized chromatin, independently of any DNA damage, thereby antagonizing 53BP1 upon formation of DSBs. The retention of the BRCA1/BARD1 complex at DSBs has been proposed to rely, at least in part, on the recruitment of HP1 proteins to DSBs. While the requirement for H3K9 methylation remains a topic of debate (Luijsterburg et al., 2009, Zarebski et al., 2009), recent evidence have shown that BARD1 can directly bind to HP1 through a PxVxL motif present in its BRCT domain, and this process retains BRCA1 at DSBs while allowing the initiation of DNA end resection (Wu et al., 2015). However, the underlying mechanism regulating this step remained unknown. In this study, we identified POGZ as an important regulator of the HP1-BRCA1-BARD1 axis and a novel player in homology-directed DNA repair pathways (Fig.6I). Importantly, our data may reconcile the two conflicting models proposed for HP1 in the repair of DSBs and POGZ may be the missing factor promoting the methylation-independent recruitment of HP1 at DNA damage sites while allowing methyltransferases (e.g. EHMT1/EHMT2), two known interactors of POGZ (Suliman-Lavie et al., 2020), to alter local methylation pattern and retain the HP1-BARD1-BRCA1 complex at DSBs.

POGZ (also called ZNF280E) has been initially proposed to promote genome stability during mitosis, by interacting with HP1-α, thus facilitating its ejection from chromosomes, a necessary step for their accurate segregation (Nozawa et al., 2010). The identification of POGZ as a partner of the adaptor protein REV7/MAD2L2 (Vermeulen, Eberl et al., 2010), a key player in the faithful seggregation of chromosomes, was initially proposed to participate in the mitotic-related role of POGZ. Interestingly, the authors noted that POGZ depletion had limited impact on the steady-state localization of HP1-α in interphase (Nozawa et al., 2010). Subsequent analysis of the role of POGZ during interphase showed that its depletion correlates with a defect in the phosphorylation of replication protein A (RPA; Ser4/Ser8) upon camptothecin treatment (Baude et al., 2016), a late marker of DNA end resection. The confirmation that POGZ is part of the proximal interactome of REV7/MAD2L2 (Findlay, Heath et al., 2018), with direct relevance for DNA repair pathway choice, suggested a possible contribution of this zinc finger protein during the DNA damage response. Still, whether POGZ may directly impact DNA repair remained, until now, unknown. The recent identification of ZNF280C/ZPET, another member of the ZNF280 subfamily, in the proximal interactome of RAD18 during genotoxic stress (Moquin, Genois et al., 2019), and its characterization as a novel inhibitor of DNA end resection further argued for a role of POGZ/ZNF280E during the repair of DSBs.

Here, we elucidated the role of POGZ in homology-directed DNA repair during interphase. Notably, our findings recapitulated several key features of HP1 during the DNA damage response. For instance, HP1 proteins dynamically accumulate and subsequently dissociate from DNA damage sites (Ayoub et al., 2008, Ayoub et al., 2009, Baldeyron et al., 2011, Luijsterburg et al., 2009, Zarebski et al., 2009), alike what we observed with POGZ, which can be detected within minutes at laser micro-irradiation before disappearing less than an hour after induction of DNA damage. Moreover, targeting HP1 has been shown to impair HR, thereby leading to a sustained G2 DNA damage checkpoint (Lee et al., 2013), resembling what we noted in POGZ-depleted cells. Importantly, we did not observe any substantial impact on the phosphorylation status and dynamics of KAP1 in POGZ-depleted RPE1-hTERT cells, unlike what has been reported with HP1 (Baldeyron et al., 2011, Goodarzi, Noon et al., 2008, White, Rafalska-Metcalf et al., 2012), likely reflecting the non-redundant contribution of the different HP1 isoforms (Bosch-Presegue, Raurell-Vila et al., 2017). Along the same lines, loss of POGZ preferentially impaired the recruitment of HP1-γ to DNA damage sites, corroborating our BioID data where we observed an increased presence of POGZ in the vicinity of HP1-γ under genotoxic stress conditions. Importantly, POGZ depletion correlated with a drastic reduction of both BRCA1 and BARD1 at DSBs, confirming previous reports linking HP1-γ and the BRCA1/BARD1 complex to DNA repair by HR (Lee et al., 2013, Wu et al., 2015, Wu et al., 2016).

Several groups have recently endeavored to better understand the role of POGZ *in vivo* (Gudmundsdottir et al., 2018, Matsumura et al., 2020, Suliman-Lavie et al., 2020). *Pogz* ablation by conventional gene targeting has highlighted its essentiality for normal murine embryogenesis (Gudmundsdottir et al., 2018), a phenotype that we confirmed in our CRISPR-mediated mouse model. Interestingly, conditional ablation of *Pogz* led to transcriptional dysregulation in hematopoietic, neural and embryonic stem cells (Gudmundsdottir et al., 2018, Suliman-Lavie et al., 2020). We tested whether POGZ could influence the DNA damage response at the transcriptional level; however, our targeted expression analysis suggests that POGZ depletion has limited impact on the expression of established HR factors. Systematic transcriptomic analysis noted that *Brca2* is downregulated in *Pogz*-/-murine fetal liver cells compared to wild-type controls (∼1.8 fold change) (Gudmundsdottir et al., 2018), an observation that we did not witness in RPE1-hTERT and HeLa cells. While it is possible that POGZ regulates DNA repair at multiple levels, including transcriptional repression, its rapid accumulation at laser micro-irradiation argues for a more direct contribution during DNA repair. Importantly, our data re-enforce the tight interdependency between POGZ and HP1 protein, highlighted by our structure-function analysis that confirmed the reliance for POGZ on its HP1-binding site for its nuclear accumulation (Matsumura et al., 2020).

The recent identification of *de novo POGZ* mutations in patients affected by a rare neurocognitive disorder (Dentici et al., 2017, Ferretti et al., 2019, Fukai et al., 2015, Matsumura et al., 2016, Samanta et al., 2020, Tan et al., 2016, Ye et al., 2015, Zhao et al., 2019a, Zhao et al., 2019b), further demonstrated the central role of this zinc finger protein *in vivo*. Interestingly, the characterization of a patient-derived mutation of POGZ in a mouse model recapitulated the main clinical features observed in the WHSUS and correlated with transcriptional dysregulation (Matsumura et al., 2020). Our *in vivo* data suggest a more complex framework, where its impact on DNA repair may contribute to the clinical features observed in WHSUS patients, suggesting that the WHSUS may be multi-factorial, with a potential “genome instability” component.

## MATERIALS AND METHODS

### Cell Lines and Transfection

HEK293T, RPE1-hTERT, and HeLa cells were cultured in Dulbecco’s Modified Eagle medium (DMEM; Wisent) supplemented with 10 % fetal bovine serum (FBS, Sigma) and 1% Penicillin-Streptomycin (P/S, Wisent). U2OS cells were cultured in McCoy’s 5A Modified medium (Wisent) supplemented with 10% FBS and 1% P/S. Primary murine B cells were maintained in Roswell Park Memorial Institute (RPMI) 1640 medium (Wisent) supplemented with 10% FBS, 5% NCTC-109 media (Thermofisher), 50 µM 2-mercaptoethanol (Sigma) and 1% P/S. All cell lines were regularly tested for mycoplasma contamination and STR DNA authenticated. Plasmid transfections were carried out using Lipofectamine 2000 Transfection Reagent (Invitrogen) following the manufacturer’s protocol. Lentivirus production was done in HEK293T by co-transfection of sgRNA or shRNA constructs, envelope protein VSV-g, and packaging plasmid psPAX2. Supernatants harvested 48-and 72-hours post-transfection and concentrated whenever possible. The U2OS cell line stably expressing an inducible mCherry-LacR-FokI was a gift of R. Greenberg (University of Pennsylvania). The DNA-repair reporter cell lines DR-GFP and SA-GFP were a gift of Dr. Jeremy Stark (City of Hope National Medical Center). The U2OS-LacO cell line was a gift from Dr. Daniel Durocher (Lunenfeld-Tanenbaum Research Institute). HeLa-POGZ-VC and HeLa-POGZ-KO cell lines were generated using pLentiCRISPRv2-puro plasmids containing sgRNA outlined in Table 2B. Cell lines were selected with 2ug/mL puromycin (Tocris) 48 hours after infection with lentivirus. Cell lines were derived from single cells, expanded and screened by immunoblotting.

### RNA Interference

All siRNAs employed in this study were siGENOME Human siRNAs purchased from Dharmacon (Horizon Discovery). RNAi transfections were performed using Lipofectamine RNAiMax (Invitrogen) using forward transfections. Except when stated otherwise, siRNAs were transfected 48 h prior to experimental procedures. The individual siRNA duplexes used are: siCTRL, D-001810-03; POGZ, D-006953-01,-18; CTIP, M-011376-00; RAD51, M-003530-04; HP1-β, M-009716-00; HP1-γ, M-010033-01; HP1-α, M-004296-02. The pLKO-puro shRNA plasmids from against POGZ (TRCN0000005707-11), HP1-α (TRCN0000062240/1), HP1-β (TRCN0000062222/3), HP1-γ (TRCN0000021916/7) were obtained from the McGill Platform for Cellular Perturbation (MPCP) as part of the MISSION^®^ shRNA library (RNAi Consortium, Broad Institute) and the nontargeting control was a gift from Dr. Marc Fabian (McGill University).

### Plasmids

The cDNAs of human POGZ, HP1-α, HP1-β, HP1-γ were obtained from the McGill Platform for Cellular Perturbation (MPCP) as part of the MISSION^®^ TRC3 human ORF collection. Quikchange site directed mutagenesis (Agilent) was performed using primers (listed in Table 2C) as per manufacturers guidelines to obtain selected POGZ mutants, furthermore, all mutant plasmids were transiently transfected into the HeLa cell line and validated by western blot. All constructs were transferred from pENTR vectors into pDEST-based constructs using LR Clonase II according to manufacturer’s instructions (ThermoFisher, 11791020). The pDEST-CMV-N-mCherry was a gift from Robin Ketteler (Addgene plasmid # 123215) (Agrotis, Pengo et al., 2019). The pDEST-mCherry-NLS-LacR plasmid was a gift from Dr. Daniel Durocher (Lunenfeld-Tanenbaum Research Institute). The pDEST-pcDNA5-BirA-FLAG N-term plasmid was a gift from Anne-Claude Gingras (Lunenfeld-Tanenbaum Research Institute). For laser micro-irradiation experiments, pDONR221 POGZ was LR recombined into a pHAGE EF1α 3xHA-tag destination vector. Plasmids encoding, I-SceI or pDEST-FRT-FLAG for the different GFP reporter assays, were kindly provided by Dr. Daniel Durocher (Lunenfeld-Tanenbaum Research Institute). Oligonucleotides containing sgRNA were phosphorylated, annealed and ligated into gel-extracted BsmbI-digested linear lentiCRISPRv2. All constructs were validated by Sanger sequencing.

### Drugs

The following drugs were used in this study: phleomycin (InvivoGen, ant-ph-1); neocarzinostatin (NCS; Sigma-Aldrich); cisplatin (Tocris); and talazoparib (BMN673; Selleckchem). Cells were pulsed with phleomycin (50 μg/ml) or NCS (100 μg/mL) for 1 hour, unless otherwise indicated, and replaced with fresh media and recovered for indicated times. Similarly, cisplatin and talazopirib were treated for 16 and 24 hours, unless otherwise indicated.

### Immunoblotting

Selected cell lines were treated as indicated prior to trypsinization, collection and PBS washes. Cells were placed in 1x LDS loading buffer (10 mM Tris–HCl, 140 mM Tris-base, 0.5 mM EDTA, 1% lithium dodecyl sulfate, 10% glycerol) with 1X protease (Roche) and phosphatase (Sigma) inhibitors. Following sonication, cell lysates were cleared by centrifugation at maximum speed for 15 min at 4°C. After the addition of loading dye and 2-mercaptoethanol, cleared lysate was placed at 70°C for 10 mins. Protein lysates were subjected to immunoblotting as previously described (Findlay, Heath et al., 2018). For phospho-protein western blots, RPE1-hTERT cells were irradiated with 1 Gy and recovered for indicated time periods. Proteins from cell lysates were obtained as previously described (Loignon, Amrein et al., 2007, Xu, Loignon et al., 2005). Protein concentration were determined using the PierceTM Micro BCA protein assay kit (ThermoFisher). 35 μg of protein was resolved on 4-12% polyacrylamide gradient Criterion XT Bis-Tris Precast Gels (Biorad Laboratories) and transferred to a nitrocellulose membrane (Sigma). Membranes were blocked with BSA 5% in Tween 20 (0.015%)-TBS for 3 hours at 4°C and probed overnight with primary antibody at a dilution 1:1000 in T-TBS. Secondary antibodies were used at a dilution of 1:10000 in T-TBS. Signal was detected using Immobilon Western Chemiluminescent HRP substrate (GE Healthcare) and X-ray films (Progene). Antibodies used are outlined in Table 1.

**Table 1.** Antibodies used for Immunoblotting, Immunofluorescence, Flow Cytometry.

**Table 2.** Primers for quantitative RT-PCR

### BioID sample preparation for Mass Spectrometry

Samples were processed from HEK 293-T Flp-In cells stably expressing Flag-BirA*HP1-β, Flag-BirA*HP1-γ and Flag-BirA*HP1-α as previously described (Findlay, Heath et al, 2018). Briefly, at 70% confluency, induction of fusion protein expression was achieved by adding 1 μM tetracycline to the cells for 24 h. After induction, the media was supplemented with 50 μM biotin, together with 150 ng/ml neocarzinostatin (NCS) where indicated, for an additional 24 h. Cells were then harvested and washed twice with PBS. Pellets were subsequently resuspended in cold RIPA buffer containing: 50 mM Tris–HCl pH 7.4, 150 mM NaCl, 1 mM EDTA, 1% NP-40, 0.1% SDS, 0.5% sodium deoxycholate, 1 mM PMSF, 1 mM dithiothreitol, 1:500 Sigma-Aldrich protease inhibitor cocktail (Sigma). Cell homogenates were sonicated and 250 U benzonase was added before high speed centrifugation (12,000 rpm, 30 min). Supernatants were incubated with pre-washed streptavidin-sepharose beads (GE) at 4°C with rotation for 3 h. Beads were collected by centrifugation (2000 rpm 1 min), washed twice with RIPA buffer and three times with 50 mM ammonium bicarbonate (ABC, pH 8.2). Beads were resuspended in 50 mM ABC and on-bead digestion was achieved by adding 1 μg trypsin (Sigma-Aldrich) to the suspension for overnight incubation at 37°C with rotation. Supernatant containing peptides was collected by centrifugation and pooled with supernatants from two following washes with HPLC-grade H_2_O. Digestion was ended with the addition of formic acid to a final concentration of 5%. Samples were centrifuged (12000 rpm for 10 min), and the supernatants were dried in a SpeedVac for 3 hr at high rate. Peptides were resuspended in 5% formic acid and kept at −80°C for mass spectrometric analysis. MS processing and protein analysis were carried out as previously described (Findlay et al., 2018). Three biological replicates were performed for each condition. Spectral counts from each replicated were averaged and normalized to protein length, yielding a normalized spectral abundance factor (NSAF) (McIlwain, Mathews et al., 2012). All gene set enrichment analyses (GSEA) were performed with NSAF values and the Reactome v7.2 gene sets using GSEA 4.0.3 (UCSD and Broad Institute) (Subramanian, Tamayo et al., 2005). Enriched pathways were then visualized using the EnrichmentMap 3.3 plugin (Bader Lab, University of Toronto) for Cytoscape v8.0 (Ideker Lab, UCSD) (Merico, Isserlin et al., 2010, Shannon, Markiel et al., 2003).

### Immunofluorescence

U2OS cells were transfected at 60% confluency and re-plated onto #1.5 coverslips of 12-15mm diameter in 24 or 12 well plates, respectively. Cells were allowed to recover for 24h, followed by selected stimulations or treatments. U2OS-LacO cell lines were transfected with 1.25-2.5 μg per well of selected plasmids containing LacR-fusion proteins and recovered for 48 hours. U2OS-FokI cells were transfected with 0.5 nmoles of selected siRNA (Dharmacon) and allowed to recover for 24 hours. In selected experiments cells would be then transfected with a siRNA-resistant construct and recovered for 24 hours prior to induction with 1 μM Shield1 (Takara Bio) and 1 μM 4-hydroxytamoxifen (Sigma) for 2 hours. Coverslips were then rinsed with PBS, twice, and fixed with 2% paraformaldehyde (PFA, Thermofisher)/PBS for 20 min at RT, washed twice with PBS, followed by a 20 min fixation in 0.3% Triton-100/PBS solution. Coverslips were washed twice with PBS and blocked for one hour in 5% BSA/0.1% Triton-100/PBS (PBSA-T) at room temperature (RT). Selected primary antibodies were then incubated for one hour at RT or overnight at 4°C. Coverslips were washed twice with PBS and incubated with fluorescent secondary antibodies (Anti-Rat Goat Alexa Fluor 488 [A-11070], Anti-Mouse Goat Alexa Fluor 647, Thermofisher) in PBSA-T for one hour at RT. Coverslips were washed twice in PBS, once in ddH2O, and then mounted with Fluoromount-G (Thermofisher). Images were acquired with a LSM800 confocal microscope (Carl Zeiss AG) and analyzed as previously described (Findlay et al., 2018). Briefly, we monitored the mean fluorescence intensity (M.F.I.) of each specific protein accumulating at the mCherry foci to the background signal for the given protein within the nucleus. Each data point represents a normalized signal on a per cell basis. For the FokI system a minimum of 25 cells were counted per biological replicate, with a total of 3 biological replicates and no less than 100 cells being counted. For γ-H2AX and BRCA1 quantification, HeLa, RPE1-hTERT, and U2OS cell lines were grown on cover slips as mentioned above. These cell lines were subjected to 1 Gy of gamma-irradiation (Faxitron, MultiRad 225) and recovered for the indicated times. Prior to cell collection, cells were incubated for one hour with 10 μM 5-Ethynyl-2’-deoxyuridine (EdU). Cells were washed twice with PBS, fixed and permeabilized, as outlined above, then subjected to EdU Click-iT labeling (Thermofisher). Cells were then processed as outlined above. Images were analyzed using the open source Java ImageJ/Fiji program. Nuclei were first identified, by thresholding on DAPI fluorescence followed by analyzing particles larger than 75 μm^2^. These regions were then used to measure fluorescence of EdU signal to identify EdU+ cells. BRCA1 and γ-H2AX foci were counted using the Find Maxima feature and Measure function. Analyses are presented as a percent of cells containing EdU in a field of view containing more than 5 respective foci. A total of 5 fields of view with a total of least 200 cells analyzed per timepoint for each cell line. For immunofluorescence of γ-H2AX in primary B cells, 50000 stimulated B-cells were collected and loaded into the cytospin chambers (ThermoScientific) following the manufacturer’s instructions and centrifuged at 500 rpm for 3 min. Slides were fixed with 4% paraformaldehyde (PFA, Thermofisher)/PBS for 10 min at room temperature (RT), washed twice with PBS, followed by a 10-minute permeabilization in 0.5% Triton-100/PBS solution. Slides were washed twice with PBS, blocked for one hour in M.O.M. Mouse IgG blocking reagent (Vector Laboratories) and blocked for one hour in 1% BSA/PBS at RT, followed by incubation with primary antibodies overnight at 4°C. Slides were washed twice with PBS and incubated with fluorescent secondary antibodies and DAPI in blocking solution for 1hr at RT. Slides were washed twice in PBS and once in ddH2O, and then mounted with Fluoromount-G (Thermofisher). Images were acquired with a LSM800 confocal microscope (Carl Zeiss AG).

### Neutral Comet Assay

A modified alkali comet assay procedure was followed as previously described (Olive & Banath, 2006). Cells were trypsinized at the indicated time points post radiation and resuspended at 30000 cells/mL in PBS. Cells were combined with low meting agarose (1%) (Sigma) at 1:3 ratio and spread over the CometSlide (Trevigen). Slides were dried at room temperature for 2 minutes and immersed into neutral lysis buffer overnight at 4°C. The next day the slides were immersed into neutral electrophoresis buffer (two 15-minute washes) followed by electrophoresis for 20 minutes. Subsequently the slides were washed in water, followed by 5-minutes incubation in 70% ethanol. Slides were dried and stained with SYBR Gold (Invitrogen). Images were acquired with a LSM800 confocal microscope (Carl Zeiss AG) and the tail moments were quantified using ImageJ with the OpenComet plugin (Gyori, Venkatachalam et al., 2014). For each condition at least 75 cells were analyzed.

### Laser micro-irradiation

U2OS stable cell populations expressing the POGZ HA-tagged was transferred to a 96-well plate with 170 μm glass bottom (Ibidi), presensitized with 10 μg/ml Hoescht 33342, and micro-irradiated using a FV-3000 Olympus confocal microscope equipped with a 405 nm laser line as described previously (Gaudreau-Lapierre, Garneau et al., 2018). Immunofluorescence was performed as described previously (Gaudreau-Lapierre et al., 2018). Briefly, following micro-irradiation, cells were allowed to recover before pre-extraction in PBS containing 0.5% Triton X-100 on ice for 15 min. Following washes with PBS, cells were fixed for 15 min in 3% paraformaldehyde, 2% sucrose PBS solution, permeabilized in PBS containing 0.5% Triton X-100 for 15 min, blocked in PBS containing 3% BSA and 0.05% Tween-20, and stained with the primary antibodies. After extensive washing, samples were incubated with 1:250 each of goat anti-mouse Alexa 488-conjugated and goat anti-rabbit Alexa 647-conjugated antibodies. DAPI staining was performed, and samples were imaged on a FV-3000 Olympus confocal microscope.

### BrdU Cell Cycle Flow Cytometry

HeLa, RPE1-hTERT, or U2OS cell lines were sub-cultured to 60% confluency. Cell lines were irradiated with 1 Gy and recovered for indicated times. Prior to cell collection, cells were incubated for one hour with 10 μM 5-bromo-2’-deoxyuridine (BrdU). Cells were harvested and immediately processed with the Biolegend Phase-Flow Alexa Fluor 647 kit (BioLegend). Cells were counterstained with DAPI and at least 10 000 events were acquired on a BD Fortessa (Becton Dickinson).

### Phosphorylation-coupled Flow Cytometry

U2OS cells were cultured until 60% confluency and, unless otherwise stated, pulsed with phleomycin were recovered for selected durations to assess recovery from DNA damage. To assess levels of phosphorylated histones or selected kinases, cells were prepared as previously described (Wu, Jin et al., 2010). Cells were collected via trypsinization and washed twice in PBS. Cells were fixed at RT in 2% PFA/PBS, followed by two washes in PBS, and then permeabilized via the addition of 95% ethanol dropwise, while vortexing, to a final concentration of 70% and stored at -20 °C until use. Cells were centrifuged at 900g and washed once with cold PBS, pelleted at 500g, and washed once with PBSA-T at RT. Cells were incubated in PBSA-T with indicated primary antibodies or isotype controls for one hour at room temperature. Cells were washed with PBS, and incubated with selected secondary antibodies for one hour at RT. Cells were washed with PBS, followed by resuspension in a PBSA-T solution containing propidium iodide (20 μg/mL, Sigma) and RNAse A (250 μg/mL, BioShop Canada). A minimum of 10,000 events were collected on a BD Fortessa (Becton Dickinson). Antibodies used are outlined in Table 1.

### Drug Cytotoxicity Screening by Sulforhodamine B Assay

HeLa, RPE1-hTERT, or U2OS cell lines were plated at a concentration of 1000 cells/well in 96 well plates. Cells were treated with indicated drugs, washed with PBS and recovered in normal medium for the remaining duration of the experiment. Cells were left to grow for 4 days and then washed twice in PBS, followed by fixation with a 10% trichloroacetic acid (TCA; Bioshop Canada Inc) at 4°C for one hour. Cells were washed in room temperature water and left to air-dry overnight. Plates were incubated in 0.04% sulforhodamine B (SRB; Sigma-Aldrich) for 30 minutes at room temperature, and washed 3 times with 0.1% acetic acid and left to air-dry. Dyed protein content was solubilized in 10mM unbuffered Tris and slight agitation at room temperature for 30 minutes. Optical density of absorbance at 530nm was acquired with a FLUOstar Optima plate reader. Control wells were used for background subtraction. Treatments were done in triplicate, averaged and normalized to an untreated control. At least 3 biological replicates were done for each drug.

### GFP-based DNA Repair Assays

For DR-and SA-GFP reporter assays, U2OS cells carrying the respective GFP expression cassette were transfected with the indicated siRNAs. Twenty-four hours after transfection, cells were transfected with empty vector (EV, pDEST-FRT-FLAG) or I-SceI plasmids. After 48 hours, cells were trypsinized, harvested, washed and resuspended in PBS. The percentage of GFP-positive cells were determined by flow cytometry. The data was analyzed using the FlowJo software and presented as previously described (Findlay et al., 2018).

### Micronuclei Formation

U2OS cells were transfected and plated on coverslips as previously described. Transfected cells were then subjected to a sublethal pulse of phleomycin (10 μg/mL) for one hour. Treated cells were cultured for 48 hours until being fixed, permeabilized and stained with DAPI before images were acquired via immunofluorescence microscopy, as previously described (Harding, Benci et al., 2017), to identify DNA containing nuclear and extranuclear bodies.

### Quantitative Real Time PCR

RNA was extracted using the RNeasy Mini kit (Qiagen). One μg of RNA was used to prepare cDNA using the *LunaScript RT SuperMix* (New England Biolabs). cDNA was then diluted 10-fold and 1 μL was used per qRT-PCR reaction. Reactions were performed in triplicate with the Luna Universal qPCR Master mix (New England Biolabs) in a total volume of 10 μL. Primers for reactions are outlined in Table 2a.

### *Pogz* Germline Mutant Mice

*Pogz* was targeted in ES cells obtained from C57BL/6J female mice using the dual-guided CRISPR/Cas9 technology. Exons 9 (ENSMUSE00001286300) and 10 (ENSMUSE00001225367) were identified as being present in all potential coding transcripts of *Pogz* and gRNAs flanking these critical exons were designed by the Toronto Centre for Phenogenomics (TCP, University of Toronto) (sgRNA sequences listed in Table 2B). These procedures involving animals were performed in compliance with the Animals for Research Act of Ontario and the Guidelines of the Canadian Council on Animal Care. The TCP Animal Care committee reviewed and approved all these procedures conducted on animals at TCP. Founder mice were screened by PCR to assess the deletion of E9/E10 and confirmed by Sanger sequencing, backcrossed and maintained on a C57BL/6J strain background. Wild type C57BL/6J mice were obtained from Jackson Laboratories. All experimental protocols with mice were performed under ethical approval from McGill Animal Care Committee. Mice were maintained under pathogen-free conditions at the Lady Davis Institute for Medical Research Animal Care Facility in accordance with institutional guidelines. Organs, tissues, plasma were collected from 8-12-week-old mice, unless otherwise stated. Mice were genotyped at weaning age (3 weeks), if required. When indicated, age-matched mice were subjected to lethal irradiation (8.5 Gy). Post-irradiation mice were given Baytril-supplemented drinking water (Bayer DVM, 2.27 mg/ml). Mice were euthanized at an established humane endpoint.

To assess kinetics of embryonic lethality and derive embryonic fibroblasts (MEFs), timed-mating experiments between *Pogz* heterozygotes were initiated. E0.5 was designated when vaginal plugs were visible. Pregnant female mice were euthanized, and embryos were harvested from uterine horns under sterile conditions. Embryos were dissected and dissociated with 0.25% trypsin at 4°C overnight, followed by 30 minutes at 37°C. MEFs were passaged twice before use in SRB assays.

For behavioral tests, age and gender matched littermates (wild type or *Pogz^+/Δ^*) mice were transported to the TCP as per Fear Conditioning and Open Field Standard Operating Protocol. For the open field experiment, each open field arena was composed of a peripheral zone (8 cm from the edge of the arena walls) and a central zone (40% of the total surface of the arena). Mice were housed in field for 20 mins, and activity is measured by IR laser detection. Activity is analyzed for duration of time spent in either zone per 5-minute bins. For the fear condition experiments, mice were housed in an experimental environment for 120 seconds (context baseline), then exposed to a cue (tone) for 30s, then exposed to a stimulus (shock) for 2 seconds and freezing was measured. The following day the mice were left in experimental environment for 5 minutes and then removed. Mice were placed back in experimental environment, for 2 minutes (Tone baseline) and then the cue was administered for 3 minutes (tone) without stimulus, and freezing time was measured.

### *Ex vivo* B cell Class Switch Recombination

To prepare primary B cells for *ex vivo* class switch recombination, splenocytes were harvested as from 8-12-week-old mice under sterile conditions and prepared, as previously described (Orthwein, Patenaude et al., 2010). Splenocytes were layered on Ficoll-Paque (GE Healthcare) to isolate lymphocytes. CD43-splenocytes were isolated by negative selection with CD43-conjugated microbeads and LR magnetic columns (Miltenyi Biotec). Cells were labelled with 0.5 μM CFSE (Thermofisher) and plated at a concentration of 0.5×10^6^ cells/mL in medium containing 50 ng/mL mouse IL-4 (R&D Systems) and 25 μg/mL lipopolysaccharide (LPS; Sigma). Unstimulated controls were set up in parallel. At indicated time points, cells were harvested to assess IgG1 expression, phosphorylated H2AX levels, and Annexin V surface expression (BD Pharmingen). Antibodies used are outlined in Table 1.

### ELISA for Determination of Immunoglobulin levels

Sandwich ELISAs were done as previously described (Heath, Fudge et al., 2018). Briefly, sera from blood was collected via cardiac puncture from 8-12-week-old mice while fecal samples were weighed and resuspended at a ratio of 100 mg feces per ml of PBS/0.01% sodium azide/1% (v/v) 100x protease inhibitor cocktail (Sigma). 96 well EIA/RIA plates (Corning) were coated overnight at 4°C with anti-isotype-specific antibodies (BD Pharmingen) in carbonate buffer, pH 9.6 to capture IgM, IgG1, IgG2b, IgG3 or IgA. Washing was done with PBS/T (0.01% Tween-20), blocking was with PBS/1% BSA, and serum and antibodies were diluted in PBS/1% BSA. Serum dilutions were incubated in the coated wells for 2h, and bound antibodies were detected using corresponding biotinylated rat anti-mouse IgM, IgG1, IgG2b, IgG3 or IgA (BD Pharmingen). This was followed by incubation with HRP-conjugated streptavidin (1:5000; Thermo Scientific) for 1h and subsequently developed using 2,2-azino-bis (3-ethylbenzothiazoline-6-sulfonic acid) substrate (Sigma). Absorbance was measured at 405nm in a BioTek Synergy HTX multi-mode reader. Standard curves and relative serum antibody concentrations were calculated using GraphPad Prism 8 software.

### Statistics

All quantitative experiments are graphed with mean +/-SEM with data from the independent number of independent experiments in the figure legend. All data sets were tested for normal distribution by Shapiro-Wilk Test. Statistical significance was determined using the test indicated in the legend. All statistical analyses were performed in Prism v8 (Graphpad Software).

### Data Availability

The mass spectrometry raw data and associated peak lists related to the BioID analysis of the different HP1 isoforms in both control and upon NCS treatment have been deposited to the MassIVE requisitory database and assigned the reference code MSV000087687.

## Supporting information

BioID analysis of the different HP1 isoforms

GSEA analysis of the HP1 BioID

Antibody List

Oligo List

## ACKNOWLEDGMENTS

We wish to acknowledge the contribution of Lauryl Nutter, Marina Gertenstein, Julie Yuan, Joanna Joeng, Igor Vukobradovic, Ann Flenniken and the technicians at the Centre for Phenogenomics (TCP, Toronto) for the generation of the *Pogz* mice and their phenotypic characterization. We are grateful to Daniel Durocher, Frederic A. Mallette, Amelie Fradet-Turcotte, Javier M. Di Noia and Chantal Autexier for critical reading of the manuscript; to Daniel Durocher, Anne-Claude Gingras, Jeremy Stark, Michael Witcher and Roger Greenberg for plasmids and other reagents. We thank Dr. Denis Faubert and Christian Poitras from the IRCM proteomics core facility. JH, MK and SF are funded by the Cole Foundation. JH and HB hold doctoral fellowships from the Fonds de Recherche Sante Quebec (FRQS). JL and BL are recipients of a Lady Davis Institute/TD Bank Studentship award. ESC received a FRQS postdoctoral training scholarship. AO is the Canada Research Chair (Tier 2) in Genome Stability and Hematological Malignancies. JFC holds the TRANSAT Chair in Breast Cancer Research. Work in the AM laboratory was supported by a CIHR Project Grant (#376288) and an NSERC Discovery Grant (# 5026). Work in the JFC laboratory was supported by a NSERC Discovery grant (RGPIN-2016-04808). Work in the MW was supported by a CIHR Project Grant (#159759). Work in the AO laboratory was supported by a CIHR Project Grant (#376245), a Transition Grant from the Cole Foundation and an internal Operating Fund from the Sir Mortimer B. Davis Foundation of the Jewish General Hospital. “The less people know, the more stubbornly they know it” – Osho.

## AUTHOR CONTRIBUTIONS

JH designed, performed most of the experiments presented in this manuscript and analyzed the data. ESC, HB, and DG performed the MS experiments and analyzed the data under the supervision of JFC. SF performed the quantification of DNA damage in *ex vivo* stimulated B-cells. VML completed the ELISA for the different serum isotypes. EPC performed the phosphor-blots analysis on POGZ-depleted cells. JL and XC performed the mouse perfusion, isolation of the murine brain, and prepared sections of brain, and SR analyzed the data. BD and TM performed laser micro-irradiation immunofluorescence experiments and AM designed the experiments and analyzed the data. BL performed micrococcal nuclease studies and MW designed the experiments and analyzed the data. MK optimized and performed initial GFP-reporter experiments. AO conceived the study, designed the research, provided supervision with JUS and wrote the manuscript with input from all the other authors.

## EXPANDED VIEW FIGURE LEGENDS

**Expanded View Figure 1.**
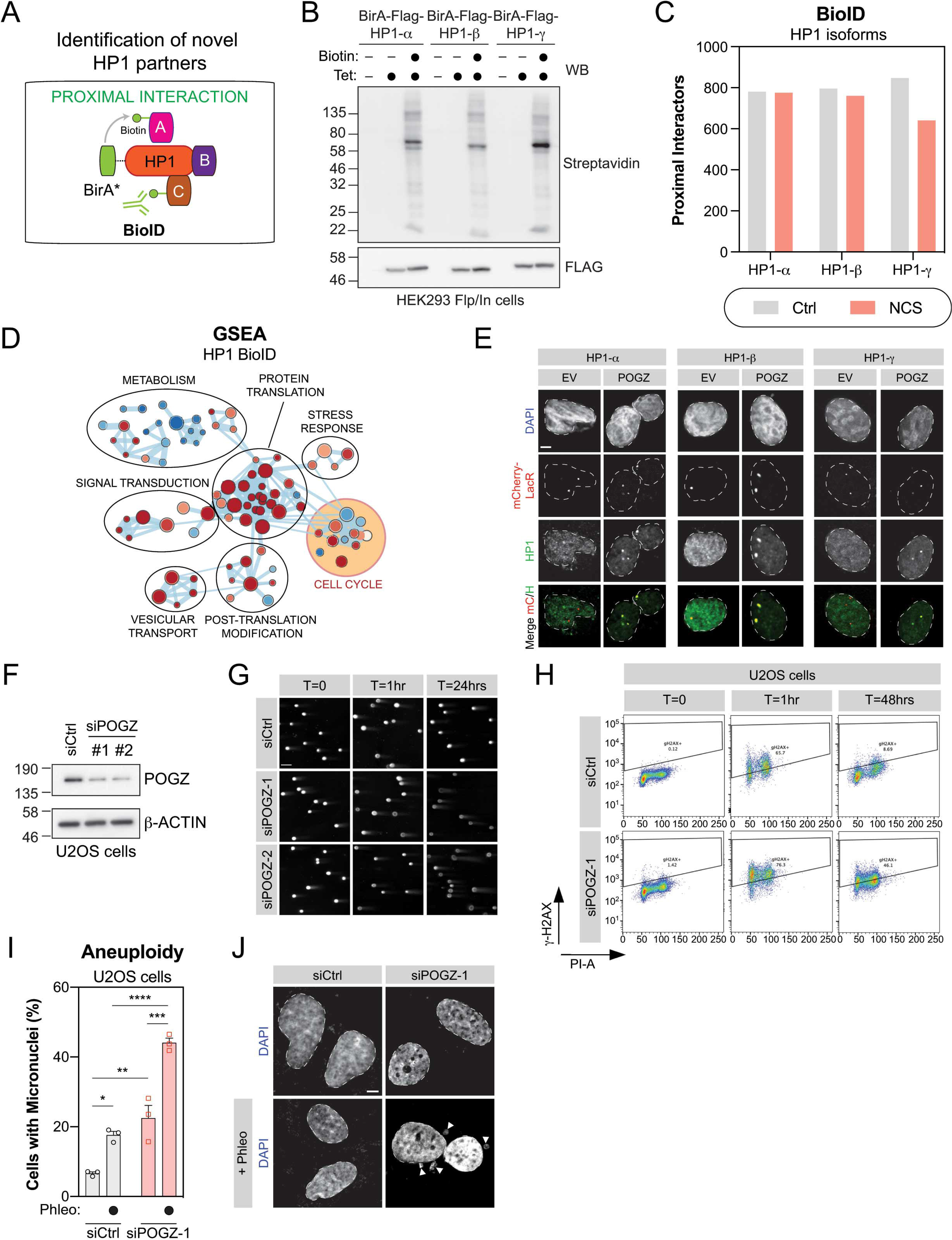
Defining the proximal interactome of the different HP1 isoforms. (A) Schematic diagram representing the BioID approach applied to HP1 and the mapping of its proximal interactome by biotinylation. (B) HEK293 Flp-In cells stably expressing each BirA*-Flag-HP1 isoform were tested for expression and biotinylation following induction with tetracycline and incubation with biotin as indicated. After induction, cells were lysed and subjected to immunoblot for Flag and Streptavidin. (C) High-confidence proximal interactors of the different HP1 isoforms identified by BioID, in presence (NCS) or absence of DNA damage (Ctrl) (n=3). (D) GSEA enrichment map of HP1-isoform interactome identifying annotated Reactome pathways. Enrichment maps from GSEA were developed with a ranked interaction network (p < 0.2, FDR < 0.5 and overlap coefficient = 0.75). Individual pathways in “Cell Cycle”, highlighted in yellow, are further examined in Fig.1C. (E) U2OS cells with a stable LacO sequence integration were transfected either with mCherry-LacR (EV) or a mCherry-LacR-tagged version of POGZ (POGZ). Immunofluorescent labeling of endogenous HP1 isoforms colocalizing with the mCherry-LacR signal were quantified and normalized to nuclear background fluorescence. Representative images of cells quantified in Fig.1E. Scale bar = 5 μm. (F) U2OS cells treated with the indicated siRNA were lysed 48hrs post-transfected and processed for POGZ western blot. β-actin was used a loading control. (G) Representative images used for quantification plotted in Fig.1F. U2OS cells treated with the indicated siRNA were irradiated, 48 hours post-transfection, with 1 Gy and run in low melting agarose under neutral conditions. DNA was stained with SYBR Gold to measure the tail moment. Scale bar = 10 μm. (H) Representative flow cytometry plots of g-H2AX levels analyzed in Fig.1G. (I) U2OS cells were transfected with indicated siRNA were stained with DAPI to visualize micronuclei by confocal microscopy. Data are number of cells per field of view displaying a micronucleus and are represented as a bar graph showing the mean ± SEM (n = 3, with a minimum of 3 fields taken per replicate). Significance was determined by two-way ANOVA followed by a Tukey’s test. \**P<0.05*, \*\**P<0.005,* \*\*\**P<0.0005,* \*\*\*\**P<0.0001*. (J) Representative images used for quantification in (H). Scale bar is 10 μm.

**Expanded View Figure 2.**
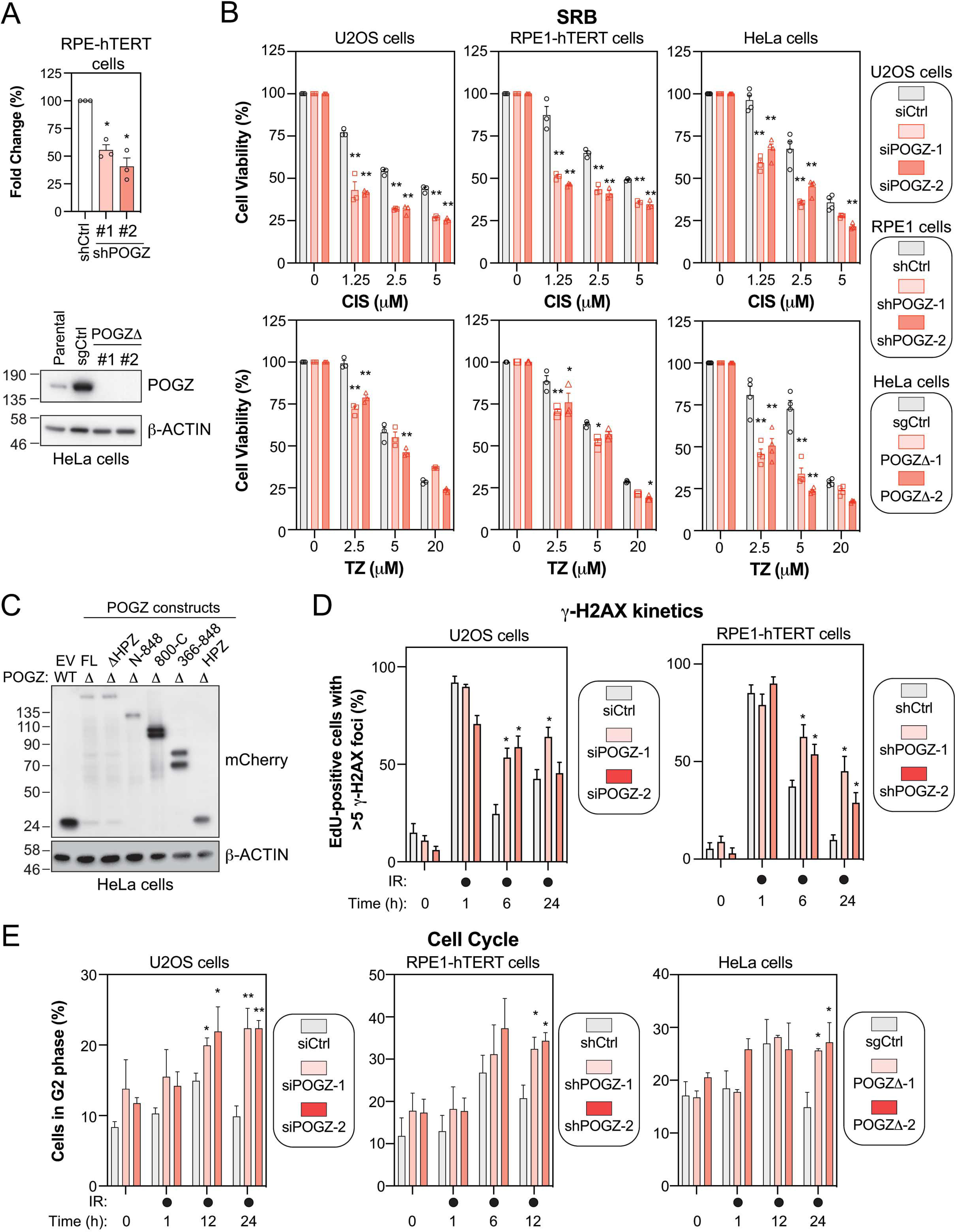
Impact of altering POGZ levels on DNA repair and cell cycle progression. (A) RPE1-hTERT cells (top panel) transduced with a scramble shRNA (shCtrl) or with a shRNA directed against POGZ (shPOGZ-1 or -2) were processed for RNA extraction. Total RNA was isolated and cDNA was generated before POGZ RNA levels were quantified by qPCR and normalized to GAPDH RNA levels. Data are represented as a graph bar showing the mean ± SEM (n=3 independent transductions for each condition). Significance was determined by one-way ANOVA followed by a Dunnett’s test. \**P<0.005.* HeLa sub-clones (bottom panel) where POGZ was targeted by CRISPR technology (POGZΔ-1 or -2) or expressing a non-targeting sgRNA control (sgCtrl), were lysed and POGZ levels were monitored by western blot. The parental HeLa cell line was added for comparison. β-Actin was used as a loading control. (B) U2OS (left panel), RPE1-hTERT (middle panel), and HeLa cells (right panel) were monitored for their sensitivities to the intercalating agent cisplatin (CIS, top panel) and the PARPi talazopirib (TZ, bottom panel) using the SRB assay. For each cell line, the following conditions were used: U2OS cells were transfected with a nontargeting siRNA (siCtrl) or an siRNA targeting human POGZ (siPOGZ-1 or -2); RPE1-hTERT cells were transduced a control shRNA (shCtrl) or a shRNA directed against human POGZ (shPOGZ-1 or -2); HeLa cells were expressing a non-targeting sgRNA (sgCtrl) or a sgRNA targeting human POGZ and sub-cloned (POGZΔ-1 or -2). Cells were pulsed with CIS or TZ at the indicated concentrations for 16 hours, or 24h, respectively, and replenished with fresh medium and incubated for 4 days. Data are represented as a bar graph showing the relative mean ± SEM, each replicate being representing as a round (Ctrl condition), square (condition #1) or triangle (condition #2) symbol. Significance was determined by two-way ANOVA followed by a Bonferroni’s test. \**P<0.05,* \*\**P<0.0001*. (C) HeLa clones where POGZ was targeted by CRISPR technology (POGZΔ) or expressing a non-targeting sgRNA control (WT), were transfected with the indicated mCherry-tagged POGZ constructs or a mCherry empty vector (EV) and lysed 48hrs post-transfection. mCherry expression was monitored by western blot. β-Actin was used as a loading control. (D) Quantification of γH2AX foci in U2OS cells transfected with the indicated siRNA (left panel), or in RPE1-hTERT cells stably expressing the indicated shRNA (right panel). Cells were treated with 1 Gy before being pulsed with Edu for 1hr and recovered at the indicated times. Data are the percentage of EdU+ cells in a field of view with >5 γ−H2AX foci and represented as a bar graph showing the relative mean ± SEM (*n* = 3, with at least 100 cells analyzed for each time point). Significance was determined by two-way ANOVA followed by a Dunnett’s test. \**P<0.005*. (E) Cell cycle distribution was monitored in U2OS (left panel), RPE1-hTERT (middle panel) and HeLa cells (right panel), transfected or transduced with the indicated condition. Cells were pulsed with BrdU for 1hr treated before being treated with 1 Gy and recovered at the indicated time points for fixation and propidium iodide staining. Data are the percentage of cells in G2 phase of the cell cycle for each indicated condition and are represented as a bar graph showing the relative mean ± SEM (n=3). Significance was determined by two-way ANOVA followed by a Dunnett’s test. \**P<0.05, **P<0.005*.

**Expanded View Figure 3.**
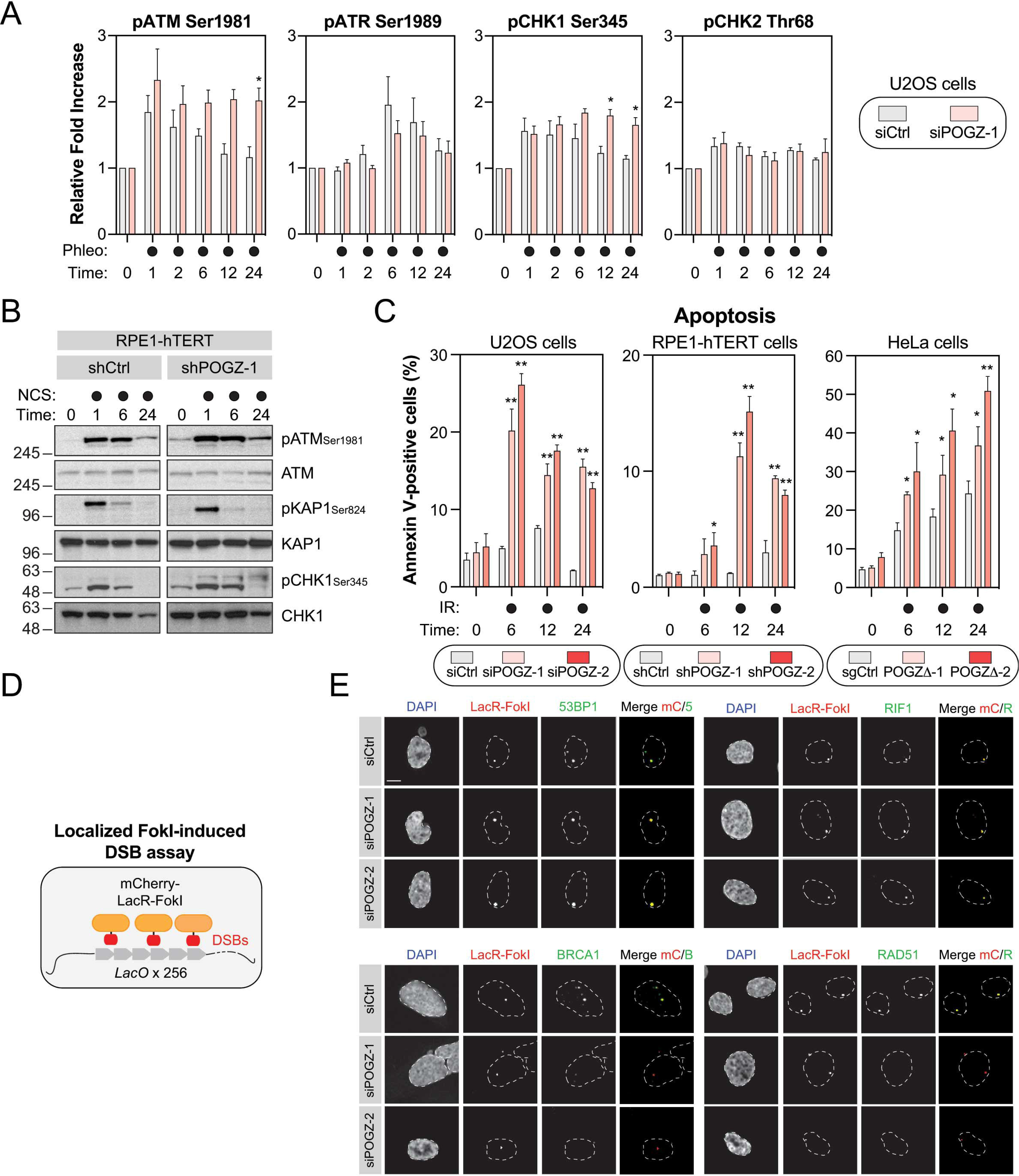
POGZ and its contribution to DNA damage checkpoint signaling and cell fate. (A) U2OS cells were transfected the indicated siRNA. 48h post-transfection, cells were pulsed with phleomycin (50 μg/mL) for 1hr before being fixed, permeabilized, and subjected to flow cytometry analysis for the indicated phospho-proteins. Data are represented as a graph bar ± SEM (n=6) where the fold increase relative to untreated samples of the respective siRNA treatment is plotted. Significance was determined by two-way ANOVA followed by a Sidak’s test. \**P<0.05*. (B) RPE1-hTERT cells transduced with the indicated shRNA were pulsed with NCS (500 μg/mL) for 1hr and harvested at the indicated time points for lysate. Western blot for the indicated phospho-and total proteins were performed and representative images are displayed in this panel. (C) Apoptosis was assessed by Annexin V staining in U2OS (left panel), RPE1-hTERT (middle panel), and HeLa cells (right panel) transfected or transduced with the indicated condition, before being treated with 10 Gy and harvested at the indicated time points for flow cytometry analysis. Data are represented as a graph bar ± SEM (n=3). Significance was determined by two-way ANOVA followed by a Dunnett’s test. \**P<0.05, **P<0.005*. (D) Schematic diagram outlining the LacR-FokI system. (E) Representative images from the U2OS-FokI data summarized in Fig.3A. U2OS-LacR-FokI cells were transfected with the indicated siRNA. The following day, DSB induction was performed with 4-OHT and Shield-1. Cells were processed for immunofluorescence microscopy using primary antibodies directed against 53BP1, RIF1, BRCA1 and RAD51. Scale bar = 2.5 μm.

**Expanded View Figure 4.**
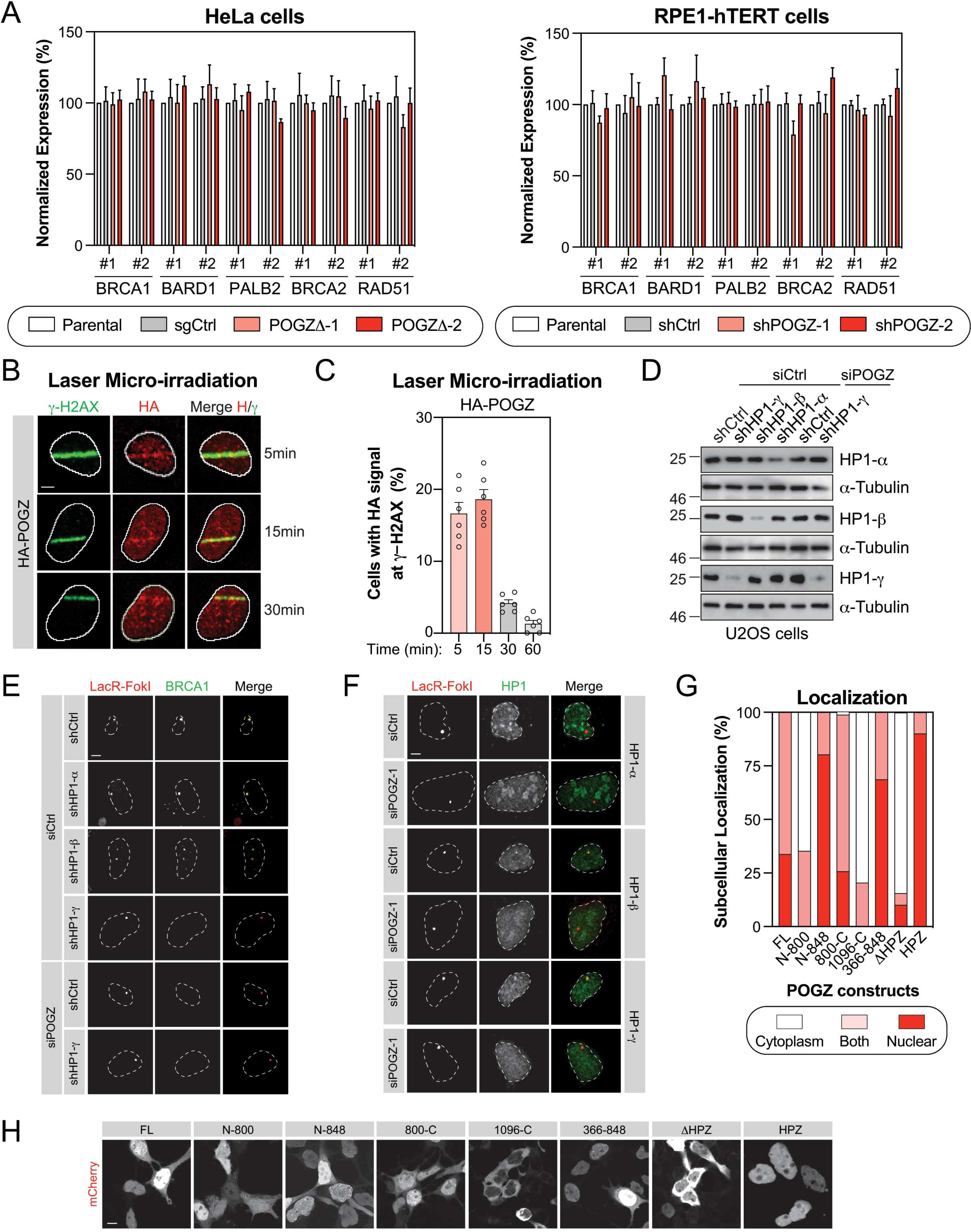
POGZ is recruited to laser micro-irradiation-induced DNA lesions. (A) RNA was extracted from HeLa cells where POGZ was targeted by CRISPR technology (left panel) or from RPE1-hTERT cells transduced with the indicated shRNA (right panel). cDNA was produced and RNA levels of the indicated HR factors were monitored by qPCR and normalized to GAPDH RNA levels. Data are represented as a graph bar ± SEM (n=3). (B) U2OS cells stably expressing a HA-tagged version of POGZ were pre-sensitized with 10 μg/ml Hoescht 33342 before being exposed to UV micro-irradiation. Immunofluorescence against HA epitope and endogenous γ-H2AX was subsequently performed to monitor POGZ accumulation at sites of DNA damage. Shown are representative micrographs of cells displaying for HA and γ-H2AX staining. Scale bar = 5 μm. (C) Quantification of U2OS cells expressing HA-POGZ and represented in (B). Data are represented as a graph bar ± SEM (n=6) where the percentage of cells with HA-POGZ signal co-localizing with γ-H2AX at the indicated time points is plotted. (D) U2OS-FokI cells stably expressing the indicated shRNA and transfected with the indicated siRNA were analyzed by western blot. 48hrs post-transfection, cells were harvested for lysate and probed for the indicated HP1 isoforms. a-Tubulin was used as a loading control. (E) Representative images of BRCA1 accumulation in U2OS-mCherry-LacR-FokI cells transduced with the indicated shRNA and transfected with the indicated siRNA. Quantification of the BRCA1 signal at the mCherry dot is represented in Fig.3E. Scale bar = 5 μm. (F) Representative images of the different HP1 isoforms accumulating at FokI-induced DSB in U2OS-mCherry-LacR-FokI cells transfected with the indicated siRNA. Quantification of the HP1 signal at the mCherry dot is represented in Fig.3G. Scale bar = 5 μm. (G) HEK293T cells were transfected with mCherry-tagged truncation constructs of POGZ or full-length mCherry-POGZ (FL). Cells were harvested 24hrs post-transfection and re-plated on to coverslips. 48 hrs post-transfection, cells were fixed, permeabilized and stained with DAPI. Cells were subsequently visualized by confocal microscopy and quantified for the presence of mCherry signal in the nucleus, the cytoplasm or both, per field of view. A minimum of 5 fields of view were sampled per biological replicate. Data are represented as a bar graph where the proportion of each sub-cellular localization is represented for each indicated construct. (H) Representative images quantified in (G). Scale bar = 5 μm.

**Expanded View Figure 5.**
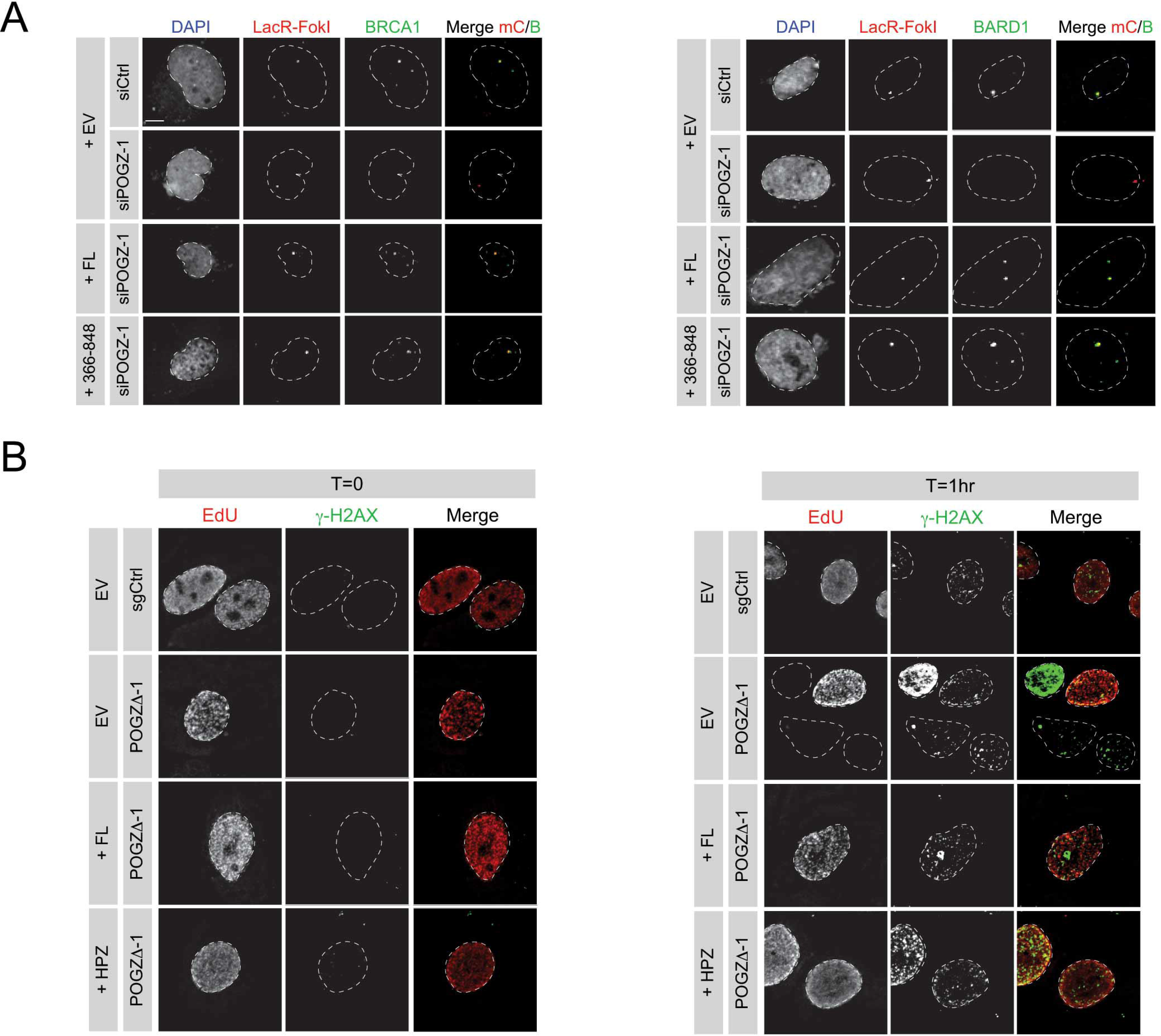
The HP1-binding site of POGZ is necessary and sufficient to restore DNA repair in POGZ-depleted cells. (A) U2OS-mCherry-LacR-FokI cells were transfected with the indicated siRNA. 24h post-transfection, cells were transfected with a Flag empty vector (EV) or a siRNA-resistant Flag-tagged POGZ construct corresponding to indicated rescue mutant. DNA damage was induced, 24h post-transfection, with Shield1 and 4-OHT. Cells were stained for BRCA1 (left panel) or BARD1 (right panel) and imaged via confocal microscopy. Representative images of the data quantified in Fig.4B. Scale bar = 5 μm. (B) HeLa cells where POGZ has been targeted by CRISPR (POGZD-1) or a control sgRNA (sgCtrl) were transfected with a mCherry empty vector (EV) or a mCherry-tagged POGZ construct corresponding to indicated rescue mutant. Cells were treated with 1 Gy and were recovered at the 1hr post-IR exposure. Cells were fixed, stained for γ-H2AX, and imaged via confocal microscopy. Representative images of the data quantified in Fig.4E. Scale bar = 5 μm.

**Expanded View Figure 6.**
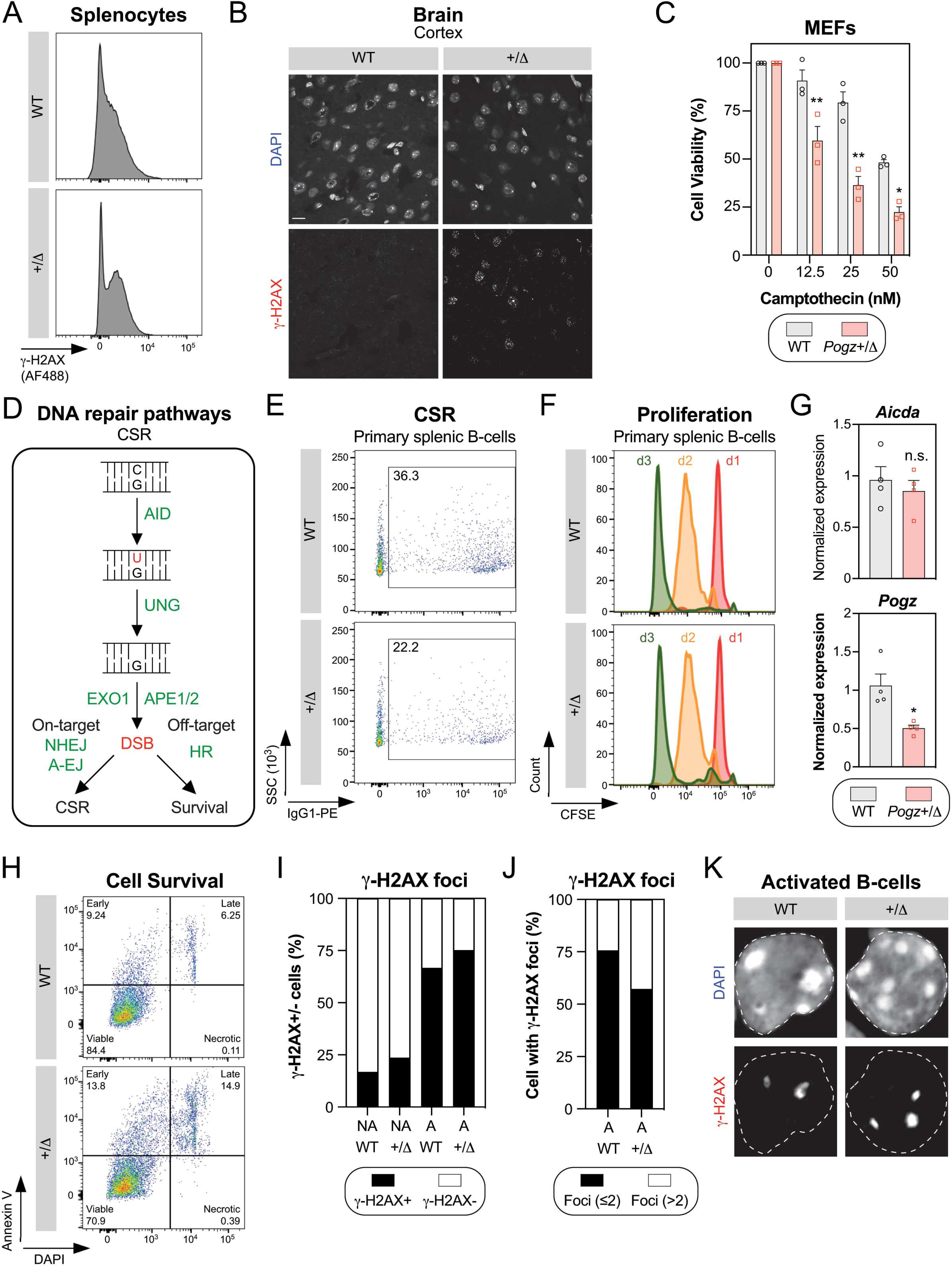
Impact of Pogz haplo-insufficiency *in vivo*. (A) Representative flow cytometry plots of γ-H2AX staining performed in splenocytes isolated from 8-week-old *Pogz* wild-type (WT) and *Pogz^+/Δ^* mice. (B) Representative micrographs of phosphorylated H2AX (γ-H2AX) from cortex tissue isolated from 6-week-old *Pogz* wild-type (WT) and *Pogz^+/Δ^* mice. Scale Bar = 20 μm. (C) MEFs obtained from the indicated genotype were treated with increasing doses of camptothecin for 1hr, before being replenished with fresh medium and incubated for 4 days. Cells were fixed and processed for SRB assay at day 4. Stained cellular content was detected by absorbance and normalized to solvent-treated conditions. Data are represented as a bar graph showing the relative mean ± SEM, each replicate being representing as a round symbol. Significance was determined by two-way ANOVA followed by a Bonferroni’s test. \**P<0.005*, \*\**P<0.0005*. (D) Schematic diagram depicting the formation and the subsequent repair of programmed DSBs in B-cells during class switch recombination (CSR) as well as the requirement for different DNA repair pathways. (E) CD43-negative primary splenocytes from 8-week-old *Pogz* wild-type (WT) and *Pogz^+/Δ^* mice were stimulated with IL-4 (50 ng/mL) and LPS (25 μg/mL). Representative flow cytometry plots monitoring surface expression of IgG1 120 hrs post-stimulation. (F) similar as in (E) except that proliferation is monitored over time by CFSE dilution. (G) 48 hrs of stimulation, total RNA was isolated from *Pogz* wild-type (WT) and *Pogz^+/Δ^* B-cells and cDNA was generated. *Aicda* and *Pogz* RNA levels were monitored by qPCR and normalized to *Gapdh*. Data are represented as mean ± SEM (n=4). Significance was determined by unpaired two-tailed t-test. \**P<0.05*. (H) similar as in (E) except that apoptosis is monitored by Annexin V/DAPI staining. (I) Stimulated CD43-negative splenocytes were loaded on to slides via Cytospin and processed for γ-H2AX immunofluorescence. Cells were fixed, stained, and imaged via confocal microscopy. Data are represented as the percentage of cells in a field of view displaying at least one γ-H2AX focus (γ-H2AX+). (J) similar as in (I) except that cells displaying at least two γ-H2AX foci (black) are represented for the indicated genotype. (K) Representative images used for the quantification plotted in (I), (J) and Fig.6H.

## REFERENCES

Agrotis A, Pengo N, Burden JJ, Ketteler R (2019) Redundancy of human ATG4 protease isoforms in autophagy and LC3/GABARAP processing revealed in cells. Autophagy 15: 976–997

Ayoub N, Jeyasekharan AD, Bernal JA, Venkitaraman AR (2008) HP1-beta mobilization promotes chromatin changes that initiate the DNA damage response. Nature 453: 682–6

Ayoub N, Jeyasekharan AD, Bernal JA, Venkitaraman AR (2009) Paving the way for H2AX phosphorylation: chromatin changes in the DNA damage response. Cell Cycle 8: 1494–500

Ayrapetov MK, Gursoy-Yuzugullu O, Xu C, Xu Y, Price BD (2014) DNA double-strand breaks promote methylation of histone H3 on lysine 9 and transient formation of repressive chromatin. Proc Natl Acad Sci U S A 111: 9169-74

Baldeyron C, Soria G, Roche D, Cook AJ, Almouzni G (2011) HP1alpha recruitment to DNA damage by p150CAF-1 promotes homologous recombination repair. J Cell Biol 193: 81–95

Barton JC, Barton JC, Bertoli LF, Acton RT (2020) Characterization of adult patients with IgG subclass deficiency and subnormal IgG2. PLoS One 15: e0240522

Bartova E, Malyskova B, Komurkova D, Legartova S, Suchankova J, Krejci J, Kozubek S (2017) Function of heterochromatin protein 1 during DNA repair. Protoplasma 254: 1233–1240

Baude A, Aaes TL, Zhai B, Al-Nakouzi N, Oo HZ, Daugaard M, Rohde M, Jaattela M (2016) Hepatoma-derived growth factor-related protein 2 promotes DNA repair by homologous recombination. Nucleic Acids Res 44: 2214–26

Bhattacharyya A, Ear US, Koller BH, Weichselbaum RR, Bishop DK (2000) The breast cancer susceptibility gene BRCA1 is required for subnuclear assembly of Rad51 and survival following treatment with the DNA cross-linking agent cisplatin. J Biol Chem 275: 23899–903

Bosch-Presegue L, Raurell-Vila H, Thackray JK, Gonzalez J, Casal C, Kane-Goldsmith N, Vizoso M, Brown JP, Gomez A, Ausio J, Zimmermann T, Esteller M, Schotta G, Singh PB, Serrano L, Vaquero A (2017) Mammalian HP1 Isoforms Have Specific Roles in Heterochromatin Structure and Organization. Cell Rep 21: 2048–2057

Botuyan MV, Lee J, Ward IM, Kim JE, Thompson JR, Chen J, Mer G (2006) Structural basis for the methylation state-specific recognition of histone H4-K20 by 53BP1 and Crb2 in DNA repair. Cell 127: 1361–73

Bunting SF, Nussenzweig A (2013) End-joining, translocations and cancer. Nat Rev Cancer 13: 443–54

Chapman JR, Barral P, Vannier JB, Borel V, Steger M, Tomas-Loba A, Sartori AA, Adams IR, Batista FD, Boulton SJ (2013) RIF1 is essential for 53BP1-dependent nonhomologous end joining and suppression of DNA double-strand break resection. Mol Cell 49: 858–71

Chapman JR, Sossick AJ, Boulton SJ, Jackson SP (2012) BRCA1-associated exclusion of 53BP1 from DNA damage sites underlies temporal control of DNA repair. J Cell Sci 125: 3529–34

Cortizas EM, Zahn A, Hajjar ME, Patenaude AM, Di Noia JM, Verdun RE (2013) Alternative end-joining and classical nonhomologous end-joining pathways repair different types of double-strand breaks during class-switch recombination. J Immunol 191: 5751–63

Densham RM, Morris JR (2019) Moving Mountains-The BRCA1 Promotion of DNA Resection. Front Mol Biosci 6: 79

Dentici ML, Niceta M, Pantaleoni F, Barresi S, Bencivenga P, Dallapiccola B, Digilio MC, Tartaglia M (2017) Expanding the phenotypic spectrum of truncating POGZ mutations: Association with CNS malformations, skeletal abnormalities, and distinctive facial dysmorphism. Am J Med Genet A 173: 1965–1969

Escribano-Diaz C, Durocher D (2013) DNA repair pathway choice--a PTIP of the hat to 53BP1. EMBO Rep 14: 665–6

Falk M, Lukasova E, Gabrielova B, Ondrej V, Kozubek S (2007) Chromatin dynamics during DSB repair. Biochim Biophys Acta 1773: 1534–45

Farmer H, McCabe N, Lord CJ, Tutt AN, Johnson DA, Richardson TB, Santarosa M, Dillon KJ, Hickson I, Knights C, Martin NM, Jackson SP, Smith GC, Ashworth A (2005) Targeting the DNA repair defect in BRCA mutant cells as a therapeutic strategy. Nature 434: 917–21

Feng L, Fong KW, Wang J, Wang W, Chen J (2013) RIF1 counteracts BRCA1-mediated end resection during DNA repair. J Biol Chem 288: 11135–43

Ferretti A, Barresi S, Trivisano M, Ciolfi A, Dentici ML, Radio FC, Vigevano F, Tartaglia M, Specchio N (2019) POGZ-related epilepsy: Case report and review of the literature. Am J Med Genet A 179: 1631–1636

Findlay S, Heath J, Luo VM, Malina A, Morin T, Coulombe Y, Djerir B, Li Z, Samiei A, Simo-Cheyou E, Karam M, Bagci H, Rahat D, Grapton D, Lavoie EG, Dove C, Khaled H, Kuasne H, Mann KK, Klein KO et al. (2018) SHLD2/FAM35A co-operates with REV7 to coordinate DNA double-strand break repair pathway choice. EMBO J 37

Fnu S, Williamson EA, De Haro LP, Brenneman M, Wray J, Shaheen M, Radhakrishnan K, Lee SH, Nickoloff JA, Hromas R (2011) Methylation of histone H3 lysine 36 enhances DNA repair by nonhomologous end-joining. Proc Natl Acad Sci U S A 108: 540–5

Fradet-Turcotte A, Canny MD, Escribano-Diaz C, Orthwein A, Leung CC, Huang H, Landry MC, Kitevski-LeBlanc J, Noordermeer SM, Sicheri F, Durocher D (2013) 53BP1 is a reader of the DNA-damage-induced H2A Lys 15 ubiquitin mark. Nature 499: 50–4

Fukai R, Hiraki Y, Yofune H, Tsurusaki Y, Nakashima M, Saitsu H, Tanaka F, Miyake N, Matsumoto N (2015) A case of autism spectrum disorder arising from a de novo missense mutation in POGZ. J Hum Genet 60: 277–9

Gaudreau-Lapierre A, Garneau D, Djerir B, Coulombe F, Morin T, Marechal A (2018) Investigation of Protein Recruitment to DNA Lesions Using 405 Nm Laser Micro-irradiation. J Vis Exp

Goodarzi AA, Noon AT, Deckbar D, Ziv Y, Shiloh Y, Lobrich M, Jeggo PA (2008) ATM signaling facilitates repair of DNA double-strand breaks associated with heterochromatin. Mol Cell 31: 167–77

Greenberg RA, Sobhian B, Pathania S, Cantor SB, Nakatani Y, Livingston DM (2006) Multifactorial contributions to an acute DNA damage response by BRCA1/BARD1-containing complexes. Genes Dev 20: 34–46

Gudmundsdottir B, Gudmundsson KO, Klarmann KD, Singh SK, Sun L, Singh S, Du Y, Coppola V, Stockwin L, Nguyen N, Tessarollo L, Thorsteinsson L, Sigurjonsson OE, Gudmundsson S, Rafnar T, Tisdale JF, Keller JR (2018) POGZ Is Required for Silencing Mouse Embryonic beta-like Hemoglobin and Human Fetal Hemoglobin Expression. Cell Rep 23: 3236–3248

Gyori BM, Venkatachalam G, Thiagarajan PS, Hsu D, Clement MV (2014) OpenComet: an automated tool for comet assay image analysis. Redox Biol 2: 457–65

Harding SM, Benci JL, Irianto J, Discher DE, Minn AJ, Greenberg RA (2017) Mitotic progression following DNA damage enables pattern recognition within micronuclei. Nature 548: 466–470

Hasham MG, Donghia NM, Coffey E, Maynard J, Snow KJ, Ames J, Wilpan RY, He Y, King BL, Mills KD (2010) Widespread genomic breaks generated by activation-induced cytidine deaminase are prevented by homologous recombination. Nat Immunol 11: 820-6

Heath JJ, Fudge NJ, Gallant ME, Grant MD (2018) Proximity of Cytomegalovirus-Specific CD8(+) T Cells to Replicative Senescence in Human Immunodeficiency Virus-Infected Individuals. Front Immunol 9: 201

Hustedt N, Durocher D (2016) The control of DNA repair by the cell cycle. Nat Cell Biol 19: 1–9

Huyen Y, Zgheib O, Ditullio RA, Jr., Gorgoulis VG, Zacharatos P, Petty TJ, Sheston EA, Mellert HS, Stavridi ES, Halazonetis TD (2004) Methylated lysine 79 of histone H3 targets 53BP1 to DNA double-strand breaks. Nature 432: 406-11

Iyama T, Wilson DM, 3rd (2013) DNA repair mechanisms in dividing and non-dividing cells. DNA Repair (Amst) 12: 620–36

Johansson P, Fasth A, Ek T, Hammarsten O (2017) Validation of a flow cytometry-based detection of gamma-H2AX, to measure DNA damage for clinical applications. Cytometry B Clin Cytom 92: 534–540

Lee YH, Kuo CY, Stark JM, Shih HM, Ann DK (2013) HP1 promotes tumor suppressor BRCA1 functions during the DNA damage response. Nucleic Acids Res 41: 5784–98

Li M, Yu X (2013) Function of BRCA1 in the DNA damage response is mediated by ADP-ribosylation. Cancer Cell 23: 693–704

Loignon M, Amrein L, Dunn M, Aloyz R (2007) XRCC3 depletion induces spontaneous DNA breaks and p53-dependent cell death. Cell Cycle 6: 606–11

Lu Y, Liu Y, Yang C (2017) Evaluating In Vitro DNA Damage Using Comet Assay. J Vis Exp

Luijsterburg MS, Dinant C, Lans H, Stap J, Wiernasz E, Lagerwerf S, Warmerdam DO, Lindh M, Brink MC, Dobrucki JW, Aten JA, Fousteri MI, Jansen G, Dantuma NP, Vermeulen W, Mullenders LH, Houtsmuller AB, Verschure PJ, van Driel R (2009) Heterochromatin protein 1 is recruited to various types of DNA damage. J Cell Biol 185: 577–86

Manke IA, Lowery DM, Nguyen A, Yaffe MB (2003) BRCT repeats as phosphopeptide-binding modules involved in protein targeting. Science 302: 636–9

Matsumura K, Nakazawa T, Nagayasu K, Gotoda-Nishimura N, Kasai A, Hayata-Takano A, Shintani N, Yamamori H, Yasuda Y, Hashimoto R, Hashimoto H (2016) De novo POGZ mutations in sporadic autism disrupt the DNA-binding activity of POGZ. J Mol Psychiatry 4: 1

Matsumura K, Seiriki K, Okada S, Nagase M, Ayabe S, Yamada I, Furuse T, Shibuya H, Yasuda Y, Yamamori H, Fujimoto M, Nagayasu K, Yamamoto K, Kitagawa K, Miura H, Gotoda-Nishimura N, Igarashi H, Hayashida M, Baba M, Kondo M et al. (2020) Pathogenic POGZ mutation causes impaired cortical development and reversible autism-like phenotypes. Nat Commun 11: 859

McIlwain S, Mathews M, Bereman MS, Rubel EW, MacCoss MJ, Noble WS (2012) Estimating relative abundances of proteins from shotgun proteomics data. BMC Bioinformatics 13: 308

Merico D, Isserlin R, Stueker O, Emili A, Bader GD (2010) Enrichment map: a network-based method for gene-set enrichment visualization and interpretation. PLoS One 5: e13984

Moquin DM, Genois MM, Zhang JM, Ouyang J, Yadav T, Buisson R, Yazinski SA, Tan J, Boukhali M, Gagne JP, Poirier GG, Lan L, Haas W, Zou L (2019) Localized protein biotinylation at DNA damage sites identifies ZPET, a repressor of homologous recombination. Genes Dev 33: 75–89

Nakamura K, Saredi G, Becker JR, Foster BM, Nguyen NV, Beyer TE, Cesa LC, Faull PA, Lukauskas S, Frimurer T, Chapman JR, Bartke T, Groth A (2019) H4K20me0 recognition by BRCA1-BARD1 directs homologous recombination to sister chromatids. Nat Cell Biol 21: 311–318

Nozawa RS, Nagao K, Masuda HT, Iwasaki O, Hirota T, Nozaki N, Kimura H, Obuse C (2010) Human POGZ modulates dissociation of HP1alpha from mitotic chromosome arms through Aurora B activation. Nat Cell Biol 12: 719–27

Olive PL, Banath JP (2006) The comet assay: a method to measure DNA damage in individual cells. Nat Protoc 1: 23–9

Orthwein A, Noordermeer SM, Wilson MD, Landry S, Enchev RI, Sherker A, Munro M, Pinder J, Salsman J, Dellaire G, Xia B, Peter M, Durocher D (2015) A mechanism for the suppression of homologous recombination in G1 cells. Nature 528: 422–6

Orthwein A, Patenaude AM, Affar el B, Lamarre A, Young JC, Di Noia JM (2010) Regulation of activation-induced deaminase stability and antibody gene diversification by Hsp90. J Exp Med 207: 2751–65

Pavone P, Pratico AD, Ruggieri M, Rizzo R, Falsaperla R (2017) Resuming the obsolete term “small head”: when microcephaly occurs without cognitive impairment. Neurol Sci 38: 1723–1725

Polo SE, Jackson SP (2011) Dynamics of DNA damage response proteins at DNA breaks: a focus on protein modifications. Genes Dev 25: 409-33

Rivera B, Di Iorio M, Frankum J, Nadaf J, Fahiminiya S, Arcand SL, Burk DL, Grapton D, Tomiak E, Hastings V, Hamel N, Wagener R, Aleynikova O, Giroux S, Hamdan FF, Dionne-Laporte A, Zogopoulos G, Rousseau F, Berghuis AM, Provencher D et al. (2017) Functionally Null RAD51D Missense Mutation Associates Strongly with Ovarian Carcinoma. Cancer Res 77: 4517–4529

Roux KJ, Kim DI, Raida M, Burke B (2012) A promiscuous biotin ligase fusion protein identifies proximal and interacting proteins in mammalian cells. J Cell Biol 196: 801–10

Samanta D, Ramakrishnaiah R, Schaefer B (2020) The neurological aspects related to POGZ mutation: case report and review of CNS malformations and epilepsy. Acta Neurol Belg 120: 447–450

Scully R, Panday A, Elango R, Willis NA (2019) DNA double-strand break repair-pathway choice in somatic mammalian cells. Nat Rev Mol Cell Biol 20: 698–714

Shanbhag NM, Greenberg RA (2013) The dynamics of DNA damage repair and transcription. Methods Mol Biol 1042: 227–35

Shannon P, Markiel A, Ozier O, Baliga NS, Wang JT, Ramage D, Amin N, Schwikowski B, Ideker T (2003) Cytoscape: a software environment for integrated models of biomolecular interaction networks. Genome Res 13: 2498–504

Stessman HAF, Willemsen MH, Fenckova M, Penn O, Hoischen A, Xiong B, Wang T, Hoekzema K, Vives L, Vogel I, Brunner HG, van der Burgt I, Ockeloen CW, Schuurs-Hoeijmakers JH, Klein Wassink-Ruiter JS, Stumpel C, Stevens SJC, Vles HS, Marcelis CM, van Bokhoven H et al. (2016) Disruption of POGZ Is Associated with Intellectual Disability and Autism Spectrum Disorders. Am J Hum Genet 98: 541–552

Subramanian A, Tamayo P, Mootha VK, Mukherjee S, Ebert BL, Gillette MA, Paulovich A, Pomeroy SL, Golub TR, Lander ES, Mesirov JP (2005) Gene set enrichment analysis: a knowledge-based approach for interpreting genome-wide expression profiles. Proc Natl Acad Sci U S A 102: 15545–50

Suliman-Lavie R, Title B, Cohen Y, Hamada N, Tal M, Tal N, Monderer-Rothkoff G, Gudmundsdottir B, Gudmundsson KO, Keller JR, Huang GJ, Nagata KI, Yarom Y, Shifman S (2020) Pogz deficiency leads to transcription dysregulation and impaired cerebellar activity underlying autism-like behavior in mice. Nat Commun 11: 5836

Tan B, Zou Y, Zhang Y, Zhang R, Ou J, Shen Y, Zhao J, Luo X, Guo J, Zeng L, Hu Y, Zheng Y, Pan Q, Liang D, Wu L (2016) A novel de novo POGZ mutation in a patient with intellectual disability. J Hum Genet 61: 357–9

Varnaite R, MacNeill SA (2016) Meet the neighbors: Mapping local protein interactomes by proximity-dependent labeling with BioID. Proteomics 16: 2503–2518

Vermeulen M, Eberl HC, Matarese F, Marks H, Denissov S, Butter F, Lee KK, Olsen JV, Hyman AA, Stunnenberg HG, Mann M (2010) Quantitative interaction proteomics and genome-wide profiling of epigenetic histone marks and their readers. Cell 142: 967–80

Vichai V, Kirtikara K (2006) Sulforhodamine B colorimetric assay for cytotoxicity screening. Nat Protoc 1: 1112–6

White D, Rafalska-Metcalf IU, Ivanov AV, Corsinotti A, Peng H, Lee SC, Trono D, Janicki SM, Rauscher FJ, 3rd (2012) The ATM substrate KAP1 controls DNA repair in heterochromatin: regulation by HP1 proteins and serine 473/824 phosphorylation. Mol Cancer Res 10: 401–14

Wu S, Jin L, Vence L, Radvanyi LG (2010) Development and application of ‘phosphoflow’ as a tool for immunomonitoring. Expert Rev Vaccines 9: 631–43

Wu W, Nishikawa H, Fukuda T, Vittal V, Asano M, Miyoshi Y, Klevit RE, Ohta T (2015) Interaction of BARD1 and HP1 Is Required for BRCA1 Retention at Sites of DNA Damage. Cancer Res 75: 1311–21

Wu W, Togashi Y, Johmura Y, Miyoshi Y, Nobuoka S, Nakanishi M, Ohta T (2016) HP1 regulates the localization of FANCJ at sites of DNA double-strand breaks. Cancer Sci 107: 1406–1415

Xu ZY, Loignon M, Han FY, Panasci L, Aloyz R (2005) Xrcc3 induces cisplatin resistance by stimulation of Rad51-related recombinational repair, S-phase checkpoint activation, and reduced apoptosis. J Pharmacol Exp Ther 314: 495–505

Ye Y, Cho MT, Retterer K, Alexander N, Ben-Omran T, Al-Mureikhi M, Cristian I, Wheeler PG, Crain C, Zand D, Weinstein V, Vernon HJ, McClellan R, Krishnamurthy V, Vitazka P, Millan F, Chung WK (2015) De novo POGZ mutations are associated with neurodevelopmental disorders and microcephaly. Cold Spring Harb Mol Case Stud 1: a000455

Young LC, McDonald DW, Hendzel MJ (2013) Kdm4b histone demethylase is a DNA damage response protein and confers a survival advantage following gamma-irradiation. J Biol Chem 288: 21376–21388

Yu X, Chini CC, He M, Mer G, Chen J (2003) The BRCT domain is a phospho-protein binding domain. Science 302: 639–42

Yuan SS, Chang HL, Hou MF, Chan TF, Kao YH, Wu YC, Su JH (2002) Neocarzinostatin induces Mre11 phosphorylation and focus formation through an ATM-and NBS1-dependent mechanism. Toxicology 177: 123–30

Zarebski M, Wiernasz E, Dobrucki JW (2009) Recruitment of heterochromatin protein 1 to DNA repair sites. Cytometry A 75: 619–25

Zhao W, Quan Y, Wu H, Han L, Bai T, Ma L, Li B, Xun G, Ou J, Zhao J, Hu Z, Guo H, Xia K (2019a) POGZ de novo missense variants in neuropsychiatric disorders. Mol Genet Genomic Med 7: e900

Zhao W, Tan J, Zhu T, Ou J, Li Y, Shen L, Wu H, Han L, Liu Y, Jia X, Bai T, Li H, Ke X, Zhao J, Zou X, Hu Z, Guo H, Xia K (2019b) Rare inherited missense variants of POGZ associate with autism risk and disrupt neuronal development. J Genet Genomics 46: 247–257

Zimmermann M, Lottersberger F, Buonomo SB, Sfeir A, de Lange T (2013) 53BP1 regulates DSB repair using Rif1 to control 5’ end resection. Science 339: 700–4

